# Legofit: Estimating Population History from Genetic Data

**DOI:** 10.1101/613067

**Authors:** Alan R. Rogers

**Affiliations:** Dept. of Anthropology, 260 Central Campus Dr., Suite 4428, Univ. of Utah, Salt Lake City, UT 84112

## Abstract

**Background:** Our current understanding of archaic admixture in humans relies on statistical methods with large biases, whose magnitudes depend on the sizes and separation times of ancestral populations. To avoid these biases, it is necessary to estimate these parameters simultaneously with those describing admixture. Genetic estimates of population histories also confront problems of statistical identifiability: different models or different combinations of parameter values may fit the data equally well. To deal with this problem, we need methods of model selection and model averaging, which are lacking from most existing software.

**Results:** The Legofit software package allows simultaneous estimation of parameters describing admixture and other aspects of population history. It includes facilities for data manipulation, estimation, model selection, and model averaging. It outperforms several statistical methods that have been widely used to study archaic admixture in humans.

## Background

Genetic data now play a prominent role in research on human prehistory. In less than a decade, we have learned that modern humans carry DNA from Neanderthal ancestors Green et al. [2010] and also from a previously unknown “Denisovan” population Reich et al. [2010], Meyer et al. [2012]; we have learned that the European Neolithic was primarily a movement of peoples Bollongino et al. [2013], Skoglund et al. [2012], but that farmers and foragers then lived side by side, exchanging genes for thousands of years Lipson et al. [2017]; we have learned that Indo-Europeans arrived in Europe about 5000 years ago as invaders from the Pontic Steppes Haak et al. [2015]; and we have learned that some populations carry DNA from “superarchaics,” which separated from other humans perhaps a million years ago Prüfer et al. [2014], Mendez et al. [2012].

There are reasons, however, to be skeptical of these new findings. First, many of the statistics used to estimate archaic admixture have large biases. For example, Rogers and Bohlender [Rogers and Bohlender, 2015, Fig. 4] document biases in one statistic that range from 50% to 600%, depending on the separation time of Neanderthals and Denisovans. Petr et al. Petr et al. [2019] show that similar bias in another statistic underlies an apparent (but artifactual) decline in the frequency of Neanderthal DNA in Europe during the past 45,000 years. To avoid these biases, one must simultaneously estimate the parameters that underlie them.

In addition to bias, there are also problems of statistical identifiability, which arise when several models fit the data equally well. Identifiability problems can lead us to prefer incorrect models of history, and they can make confidence intervals unrealistically narrow. Consequently, it is likely that some of the recent findings summarized above are incorrect.

The Legofit package Rogers et al. [2017a,b] introduces methods that address these problems. It reduces bias by allowing simultaneous estimation of the parameters that introduce bias into competing estimators. It uses model selection and model averaging to cope with identifiability problems, and it uses residual analysis to diagnose misspecified models. This article will not attempt a comprehensive review of genetic methods for estimation of population history. Instead, it will describe Legofit and compare it against several methods that are widely used in the study of archaic admixture.

## Implementation

### Nucleotide site patterns

Legofit works with the frequencies of *nucleotide site patterns*, which are defined below. The first step in any analysis involves tabulating site pattern frequencies from data. Legofit provides tools that tabulate these frequencies from standard data formats and also from several forms of simulation output.

Site patterns are illustrated in Fig. 1. A nucleotide site exhibits the *yn* site pattern if random nucleotides drawn from populations *Y* and *N* carry the derived allele, but those drawn from other populations carry the ancestral allele. They represent the special case of the site frequency spectrum Hudson [2015] in which the sample consists of one haploid genome per population.

**Figure 1:**
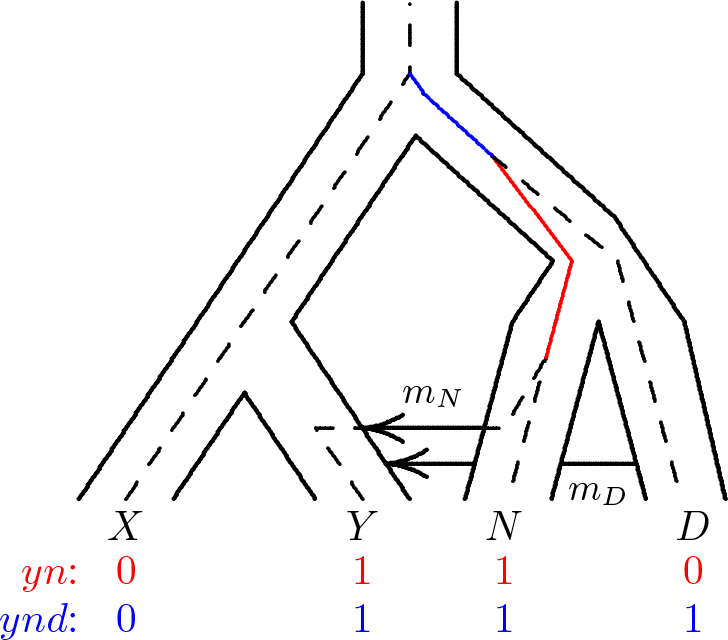
Population tree with embedded gene tree. A mutation on the solid red branch would generate site pattern *yn* (shown in red at the base of the tree). One on the solid blue branch would generate *ynd*. “0” and “1” represent the ancestral and derived alleles. Key: *X*, Africa; *Y*, Eurasia; *N*, Neanderthal; *D*, Denisovan.

In Fig. 1, a mutation on the red branch would generate *yn*, whereas one on the blue branch would generate *ynd*. Mutations elsewhere would generate other site patterns. Let *B*_*i*_ represent the length in generations of the branch generating site pattern *i*. For example, *B*_*yn*_ is the length of the red branch in Fig. 1 and *B*_*ynd*_ is the length of the blue branch. In any given gene tree, many of these lengths will be zero. For example, *B*_*xy*_ = 0 in Fig. 1, because no single mutation on that gene tree could generate site pattern *xy*.

Conditional on *B*_*i*_, the number of mutations on the branch generating pattern *i* is Poisson with mean *uB*_*i*_, where *u* is the mutation rate per nucleotide site per generation. We use the model of infinite sites Kimura [1969], which assumes that *u* is small enough that we can ignore the possibility of multiple mutations on a given branch. To this standard of approximation, the unconditional probability of site pattern *i* on a random gene tree is *uE*[*B*_*i*_], where the expectation is with respect to the coalescent process constrained by the network of populations.

Let *I*_*i*_ represent the count of site pattern *i* across all sequenced nucleotide positions. It’s expected value is *E*[*I*_*i*_] = *uLE*[*B*_*i*_], where *L* is the number of nucleotide positions in the sequence. The probability that a particular polymorphic site exhibits pattern *i* is

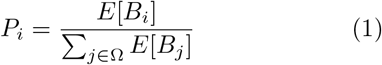

where Ω is the set of site patterns under study.

In previous publications Durand et al. [2011], Rogers and Bohlender [2015] we and others have derived analytical expressions for *E*[*B*_*i*_] under particular models of history. This analytical approach becomes difficult as models grow in complexity. Legofit relies instead on computer simulations, which make it feasible to deal with complex models of history. In each iteration of the simulation, the coalescent algorithm builds a gene genealogy analogous to the one in Fig. 1. From this genealogy, legofit calculates branch lengths (*B*_*i*_). It estimates *E*[*B*_*i*_] as the average of *B*_*i*_ across simulation replicates. Eqn. 1 then estimates *P*_*i*_.

This approach simulates branch lengths but not mutations, and the simulations can be done in parallel. For a given level of accuracy, it is orders of magnitude faster than programs that simulate both mutation and recombination. This speed makes it possible to deal with the entire suite of site patterns and with complex models involving tens of populations. We have validated it by comparison with theoretical results in models for which analytical theory is feasible Rogers and Bohlender [2015]. We can also validate by comparing the expected values generated by our method to data simulated in other ways. This is done in Fig. 2, which shows that all three simulators generate distributions of site pattern frequencies that are centered around the expected values estimated by legofit. This verifies the reliability of our approach.

**Figure 2:**
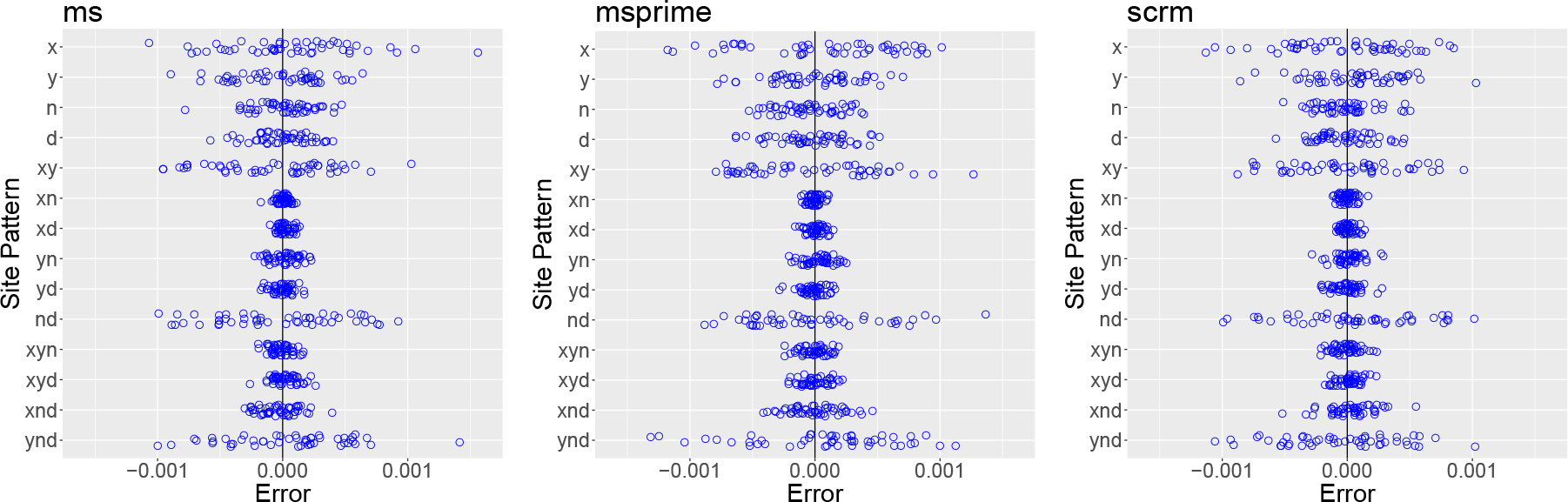
Deviation from expected values in 50 data sets generated by each of three simulation programs: ms Hudson [2002], msprime Kelleher et al. [2016], and scrm Staab et al. [2015]. All simulations assume the same model of history, which is illustrated in Fig. 1 and described fully in the additional file. Expected values were calculated with legosim. Blue circles show 50 simulated data sets.

### Models of history

A model of population history is specified in a file whose name ends with “.lgo.” This file specifies the population tree and the location of genetic samples within it. It also specifies how population size varies throughout the tree and the times at which populations separate or introgress. These parameters fall into three categories: (1) *free* parameters are estimated by legofit; (2) *fixed* parameters have values that do not change; and (3) *constrained* parameters are specified as known functions of one or more other parameters. Constrained parameters model relationships among variables that are implied either by theory or by analysis of variation among bootstrap or simulation replicates. We use them below to reexpress free variables in terms of principal components.

### Tabulating site patterns from data

The first stage of analysis involves tabulating site patterns from DNA sequence data. These data need not be phased, but they should be free of ascertainment bias. In the discussion above, I assumed that one haploid genome is sampled from each population. Real samples are larger, and a given nucleotide site may contribute to several site patterns. The contribution to a given site pattern is the probability that a sub-sample, consisting of one haploid genome drawn at random from the larger sample of each population, would exhibit this site pattern. For example, consider a model with three populations, *X*, *Y*, and *N*, and let *p*_*iX*_, *p*_*iY*_, and *p*_*iN*_ represent derived allele frequencies at the *i*th polymorphic site in the samples from these populations. Then site pattern *xy* occurs at site *i* with probability *z*_*i*_ = *p*_*iX*_*p*_*iY*_ (1 − *p*_*iN*_) [Green et al., 2010, p. S131]. Aggregating over sites, *I*_*xy*_ = ∑_*i*_z_i_ summarizes the information in the data about this site pattern. In general, for the *j*th site pattern, the analogous summary is *I*_*j*_. In this formulation *I*_*j*_ is no longer a count. It is the expected count in a random subsample of the full sample.

The Legofit package includes programs for tabulating site patterns from data and from several publicly-available programs for coalescent simulation: ms Hudson [2002], msprime Kelleher et al. [2016], and scrm Staab et al. [2015].

### Estimation

Legofit estimates parameters by maximizing the composite likelihood,

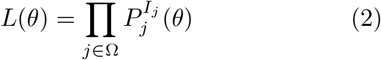

where *P*_*j*_ is as given in Eqn. 1, Ω is the set of site patterns under study, and *θ* is a vector of free parameters. This is not the full likelihood, because it ignores linkage disequilibrium and treats nucleotide sites as though they were independent.

Legofit uses a numerical algorithm—differential evolution [DE, Price et al., 2006]—to maximize *L*. DE maintains a swarm of points, which are initially distributed widely across the parameter space. In each generation, these points mutate and recombine to form offspring, which then undergo selection to form the next generation. The objective functions of the points are evaluated in parallel, in separate threads of execution. This process involves several stages, beginning with an initial stage in which the objective function is evaluated with modest precision and progressing to a final stage, which typically uses two million simulation replicates per function evaluation. This provides much more precision than a sample of two million polymorphic nucleotide sites, because we are simulating branch lengths only—not mutation or recombination.

### Bootstrap confidence intervals

The Legofit package uses a bootstrap Efron and Tibshirani [1993] to measure uncertainty. Because linked loci are not statistically independent, we cannot use an ordinary bootstrap. Instead, Legofit uses a moving-blocks bootstrap Liu and Singh [1992], which resamples blocks of nucleotides. By default, each block consists of 500 polymorphic nucleotide sites.

Bootstrap replicates approximate independent samples from the stochastic process that produced the original data. By applying legofit to many bootstrap replicates, we obtain an approximation of the sampling distribution of the estimates. This distribution is used to estimate confidence intervals.

Each bootstrap replicate is analyzed by a separate instance of the legofit program. These instances can operate in parallel, on separate nodes of a compute cluster. Legofit is thus parallel in two senses: within each node, legofit uses multiple threads to parallelize across the points maintained by the DE algorithm. It also uses multiple nodes to parallelize across bootstrap replicates.

### Model selection

The study of population history requires that we choose among complex, non-nested models. Better fits can usually be achieved with more complex models, but this improvement may be illusory—the consequence of fitting noise rather than signal. Over-fitting, as this is called, can produce incorrect inferences about population history Hawkins [2004]. We may report evidence of gene flow or of bottlenecks in population size where no such inference is warranted. Reliable inference requires that we protect against overfitting. This is not possible with the genetic methods currently used to study archaic admixture.

In other statistical contexts, such problems might be addressed via tools such as Akaike’s information criterion [AIC, Akaike, 1974] or the Bayesian information criterion [BIC, Schwarz, 1978], which penalize complex models in a principled way. These tools, however, require access to the full likelihood function, which is never available for genome-scale data sets.

Because of the size and complexity of the human nuclear genome, all statistical methods simplify the problem in some way. Legofit uses *composite likelihood*, which ignores genetic linkage and treats nucleotide sites as though they were statistically independent. This produces unbiased estimates but does not allow us to use AIC or BIC to protect against overfitting.

Legofit provides two methods of model selection: the *bootstrap estimate of predictive error* [bepe, Efron, 1983, Efron and Tibshirani, 1993], and a *composite likelihood information criterion* [clic, Varin and Vidoni, 2005].

### Bootstrap estimate of predictive error (bepe)

Bepe is analogous to cross-validation, but uses bootstrap replicates instead of partitions of the data. The first step in the process uses legofit to fit a given model to each bootstrap replicate. These runs report the predicted frequency of each nucleotide site pattern. Legofit’s “bepe” program then calculates the mean squared difference between these bootstrap-predicted frequencies and those in the real data and applies a small bias correction. The resulting estimate of predictive error compares favorably with cross-validation [Efron and Tibshirani, 1993, sec. 17.6]. It is convenient, because we need bootstraps anyway for confidence intervals.

### Composite likelihood information criterion (clic)

Clic generalizes Akaike’s information criterion [AIC, Akaike, 1974] to the case of composite likelihood. Varin and Vidoni [Varin and Vidoni, 2005, p. 523] define an information criterion that is the negative of

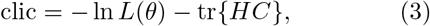

I have reversed the sign so that we can select models by minimizing (rather than maximizing) clic. In this expression, *L* is composite likelihood (Eqn. 2), *θ* is the vector of parameters, *C* is a matrix whose *ij*th entry is the sampling covariance between the *i*th and *j*th parameters, and *H* is the expectation of the negative of the Hessian matrix, and “tr” represents the matrix trace.

I estimate *C* from covariances across bootstrap or simulation replicates. *H* is a matrix of expectations of second-order partial derivatives of ln *L* with respect to pairs of parameters. Rather than taking these expectations, I evaluate the derivatives at the maximum composite likelihood estimate, 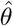 Efron and Hinkley [1978]. Within a small neighborhood near 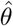, ln *L* can be approximated by a quadratic surface,

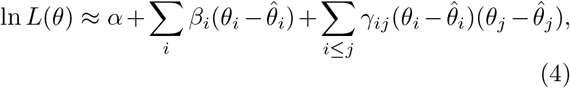

where *α* is the *Y* intercept, and *β*_*i*_ and *γ*_*ij*_ are regression coefficients.

I estimate *α*, *β*_*i*_, and *γ*_*ij*_ by ordinary least squares, using points in the neighborhood of the estimate, 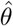. Then *H* is assembled using the second-order derivatives of ln *L*, as implied by Eqn. 4. Finally, *C* and *H* are used with Eqn. 3 to calculate clic.

### Bootstrap model averaging (booma)

Below, we will consider three models whose bepe values are 2.17 × 10^*−*7^, 5.54 × 10^*−*7^, and 6.17 × 10^*−*5^. The first model has the smallest value and is therefore preferred. But the other values are also small. Are we justified in ignoring them? To answer this question, let us consider the problem of model averaging.

When no model is clearly superior, it is better to average across several than to choose just one Buckland et al. [1997]. Otherwise, confidence intervals are misleadingly narrow because they ignore uncertainty about the model itself. In model averaging, individual models are assigned weights as discussed below. Parameters are estimated as the weighted average of estimates from individual models. Most authors rely on information criteria to provide the weights Claeskens and Hjort [2008]. One could use clic in this way, but I prefer *bootstrap model averaging* Buckland et al. [1997], which works with either bepe or clic.

This method is implemented by the Legofit program “booma.” Some model selection criterion (bepe or clic) is calculated separately for the real data and for each bootstrap replicate. (To calculate bepe for a bootstrap replicate, we pretend that the replicate is real data and the real data are a bootstrap replicate.) If there are 50 bootstrap replicates, this process gives us 51 values of the model selection criterion for each model. For each of these 51 cases, booma asks which model “wins,” i.e., which has the lowest value of the criterion. The weight of the *i*th model is the fraction of cases in which it is the winning model.

Using these weights, booma averages across models to obtain a model-averaged estimate of each parameter. If a parameter is present in only a subset of the models, the weights are re-normalized so that they sum to unity across this subset. This averaging is applied not only to the real data but also to each bootstrap replicate. This allows us to estimate confidence intervals for model-averaged estimators.

If one model is clearly superior, its weight will be unity and those of the other models will be zero. This provides a simple criterion for choosing one model over its alternatives. For the three models mentioned at the top of this section, the weights were 1, 0, and 0. This implies that the differences among the bepe values are large compared to those expected in repeated sampling from the stochastic process that generated the original data. We are therefore justified in rejecting all models but the first. This analysis is described in more detail below.

### Identifiability and principal components

Fig. 3 illustrates a problem of statistical identifiability, which arises frequently not only with Legofit, but with all methods that estimate complex population histories. Each panel in the figure is a bivariate scatterplot comparing two parameters. Each point indicates the estimated values of the two parameters in one simulation replicate. In several panels, the points fall along straight lines, indicating that the parameters are tightly correlated. These associations represent ridges in the composite likelihood surface and imply that our statistical problem has fewer dimensions than parameters. This does not lead to incorrect inferences, but it does broaden the confidence intervals of the parameters involved.

**Figure 3:**
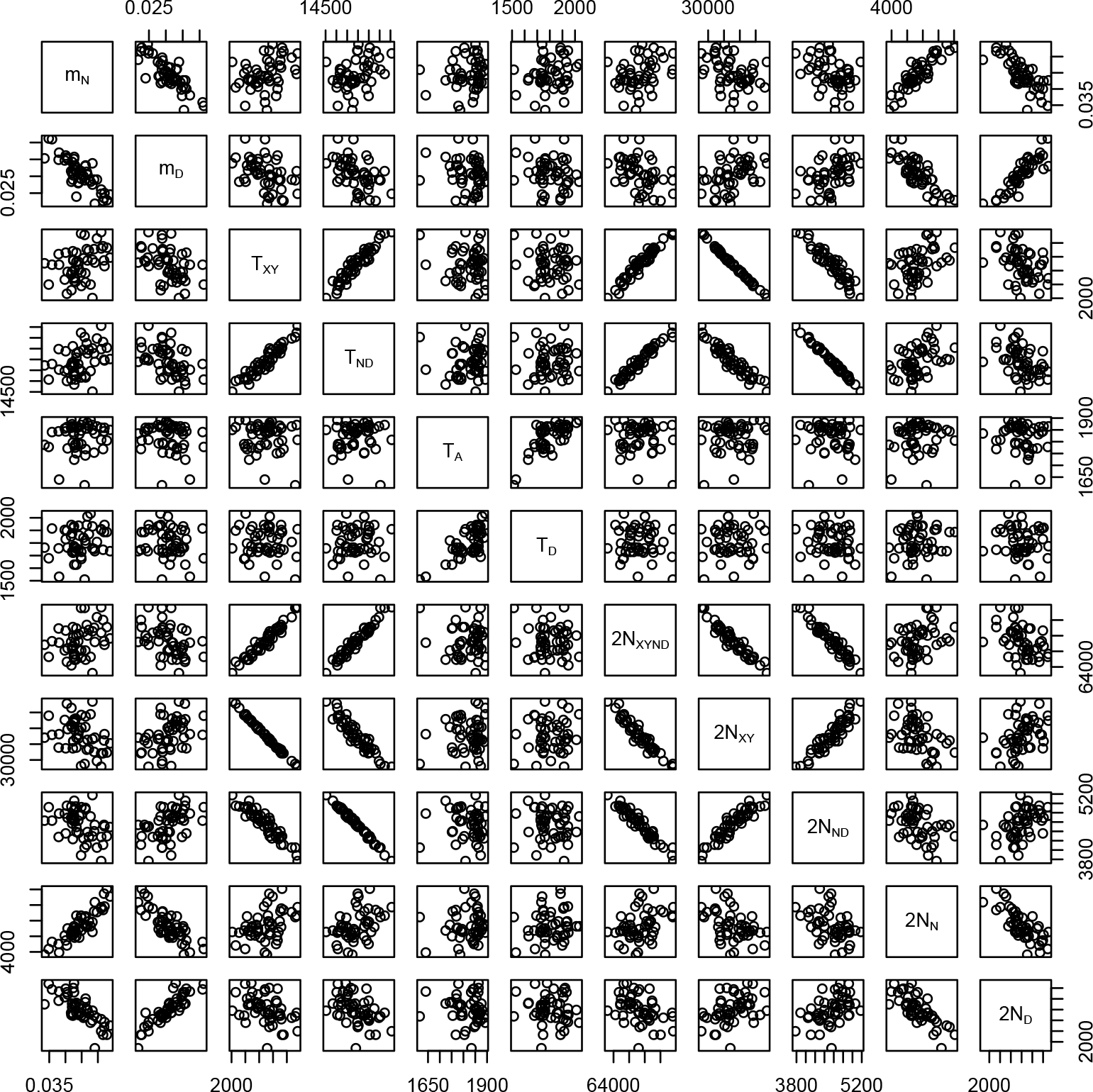
Associations between pairs of parameter estimates in 50 data sets simulated with msprime Kelleher et al. [2016] under the model in Fig. 1. Key: *m*_*N*_, fraction of admixture from *N* into *Y*; *m*_*D*_, fraction of admixture from *D* into *Y*; *T*_*XY*_, separation time of *X* and *Y*; *T*_*ND*_ separation time of *N* and *D*, *T*_*A*_, age of fossil genome from population *N*; *T*_*D*_, age of fossil from *D*; *N*_*XY* *ND*_, size of ancestral population; *N*_*XY*_, size of population ancestral to *X* and *Y*; *N*_*ND*_, size of population ancestral to *N* and *D*; *N*_*N*_, size of population *N*; *N*_*D*_, size of population *N*. The separation time, *T*_*XY* *ND*_, of *XY* and *ND* was fixed exogeneously to calibrate the molecular clock.

These problems can be ameliorated by reducing the dimension of the parameter space. The Legofit package includes pclgo, a program that calculates principal components from the bootstrap replicates and then uses these to re-express the free variables in terms of principal components. Predictive error (as measured by bepe) can be improved by excluding principal components with small eigenvalues. This usually tightens confidence intervals.

By default, pclgo merely re-expresses the free variables in terms of the principal components, and there is no reduction in dimension. To reduce dimensionality, the user must specify a tolerance criterion. The command `pclgo --tol` 0.001 would include only those components that explain at least a fraction 0.001 of the variance. Different choices of this tolerance criterion constitute different models, and we can choose among them using bepe or clic, together with booma.

## Results

Rogers and Bohlender Rogers and Bohlender [2015] document pronounced biases in the statistics that underlie our current understanding of archaic admixture. These biases are profound if there are multiple sources of admixture. To check for such bias in legofit, I simulate data under the model in Fig. 1, which allows gene flow into Eurasia (*Y*) not only from Neanderthals (*N*), but also from Denisovans (*D*). Details of this model and of all the analyses below can be found in the additional file. Here, I summarize results.

Figure 4 shows the true parameter values (red crosses) and sampling distributions (blue circles) estimated using legofit from 50 independent simulation replicates. I used pclgo to reduce dimensionality. This involves excluding dimensions that explain less than some arbitrarily-chosen fraction of the variance. I considered three models: one in terms of the original variables (without using pclgo), one using principal components with no reduction of dimension, and one excluding components that explain less than a fraction 0.001 of the variance. The weights of these three models are 0, 0.42, and 0.58 using bepe and 0, 0.12, and 0.88 using clic. Thus, pclgo seems to improve estimates, especially when some principal components are excluded. Fig. 4 shows the bepe version of the model-averaged estimates.

**Figure 4:**
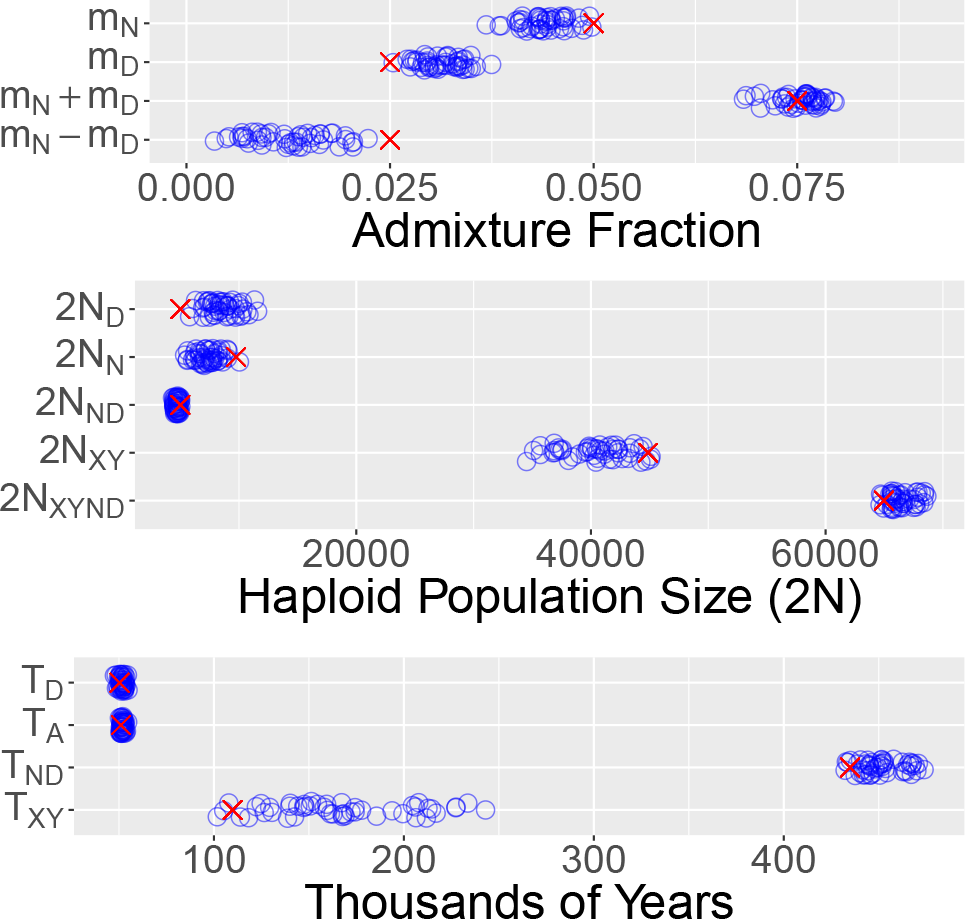
Sampling distributions of legofit estimates based on the 50 simulated data sets shown in Fig. 3. Red crosses represent true parameter values. Points have been vertically jittered to reduce overplotting in this figure and in those that follow.

All of the sampling distributions enclose the true parameter values, and several are reassuringly narrow. Nonetheless, some bias is evident in the distributions of Neanderthal admixture (*m*_*N*_) and Denisovan admixture (*m*_*D*_). The mean estimates of these parameters are closer together than are the true parameter values. This is because Neanderthals and Denisovans are sister populations, and it is hard to tell them apart. We get a better estimate of total archaic admixture, *m*_*N*_ + *m*_*D*_, than of the difference, *m*_*N*_ − *m*_*D*_.

For comparison with legofit’s estimates of the admixture fraction, Fig. 5 shows the behavior of three previously-published estimators Reich et al. [2010], Meyer et al. [2012] that have been used to study archaic admixture in humans. Nea and den work by comparing the frequencies with which derived alleles are shared by pairs of samples from different populations. Nea has also been called *R*_Neandertal_ Reich et al. [2010]. Rogers and Bohlender Rogers and Bohlender [2015] show that these estimators have large biases, especially when (as in the present model) a population receives gene flow from more than one source. Thus, it is no surprise that nea and den exhibit large biases in Fig. 5. Indeed, the black triangles show that the observed bias is in good agreement with theoretical expectations.

**Figure 5:**
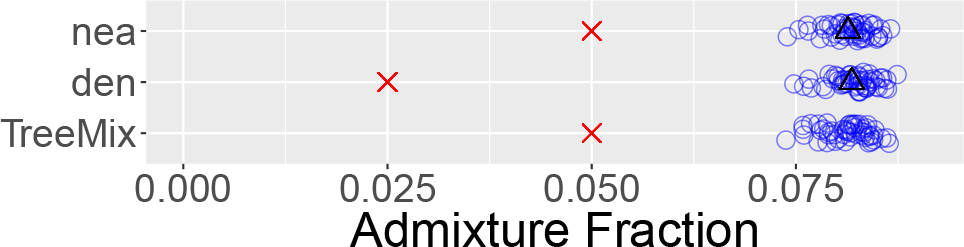
Bias in three previously-published estimators of archaic admixture. Nea and den [Meyer et al., 2012, supp. note 11] estimate Neanderthal and Denisovan admixture. TreeMix Pickrell et al. [2012] estimates Neanderthal admixture. Key: blue circles, estimates from simulated data shown in Fig. 3; red crosses, true parameter values; black triangles, expected values of statistics.

Many studies have cited an estimate that about 6% of Papuan DNA derives from Denisovans. This result is due to Meyer et al. Meyer et al. [2012], who inferred it using TreeMix Pickrell et al. [2012]. However, these authors suspected that the result was biased, because their analysis excluded Neanderthals [Meyer et al., 2012, supp. note 12]. The TreeMix results in Fig. 5 should avoid this problem, because Neanderthals are included along with Denisovans and moderns from Africa and Eurasia. TreeMix was able to detect a signal of gene flow from Neanderthals into Eurasians. As the figure shows, however, its estimate of the admixture fraction was profoundly biased. TreeMix was unable to detect gene flow from Denisovans into Eurasians. This episode of gene flow did not appear in the output from any of the simulation replicates. Instead, TreeMix reported evididence of gene flow in various parts of the tree. These episodes of gene flow were not consistent from replicate to replicate and did not exist in the simulation model.

In Fig. 4, we had the advantage of working with the true model of history. This is never the case with real data. Let us therefore consider how the analysis might proceed if we did not know the true model in advance. We would start by examining site pattern frequencies, which are shown in Fig. 6. The most common patterns (apart from singletons) are *xy* and *nd*, reflecting the shared ancestry of populations *X* and *Y* and of *N* and *D*. Let us therefore fit a model with a tree of form ((*X, Y*), (*N, D*)). This model is misspecified, because it omits gene flow. The residuals of this model are shown in Fig. 7 along with those of a correctly-specified model. The misspecified model generates many residuals that are far from zero, and these discrepancies provide clues about what is wrong with the model. For example, note that the misspecified model has positive residuals for *yn* and *ynd* but a negative residual for *y*. This suggests that we should add *N* → *Y* gene flow to the model, because such gene flow inflates the first two of these site patterns but deflates the third.

**Figure 6:**
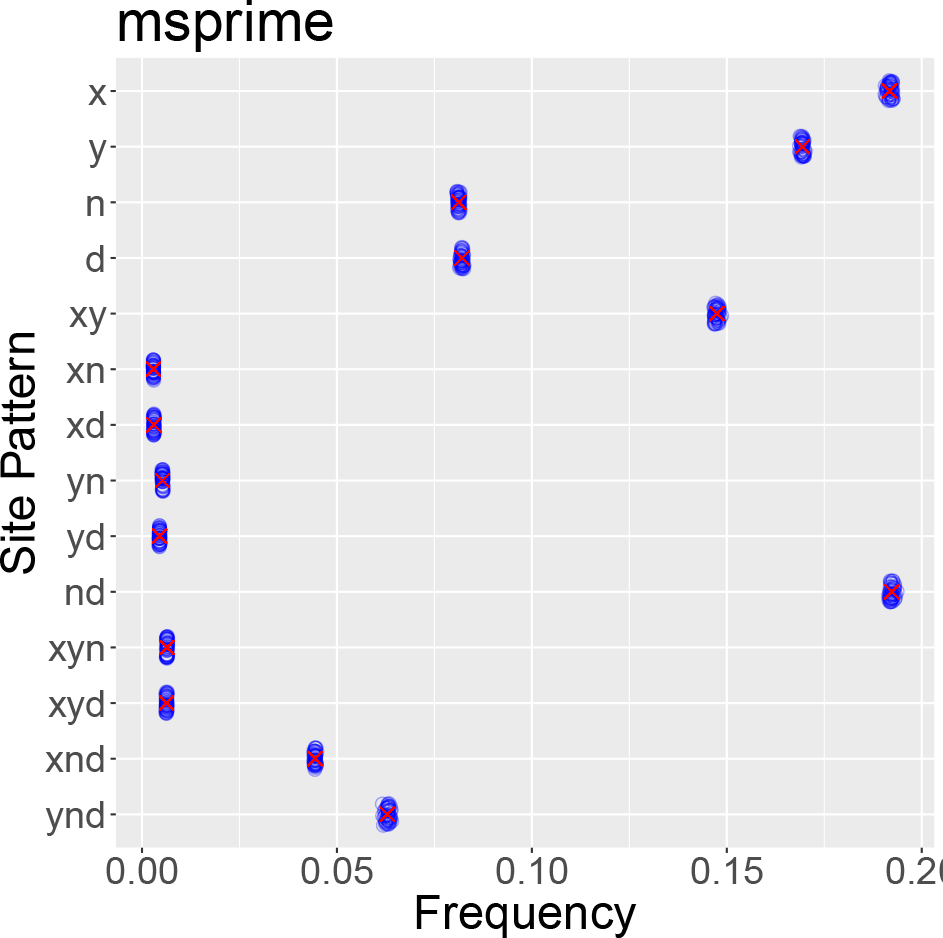
Site pattern frequences simulated using msprime Kelleher et al. [2016] under the model in Fig. 1. Data are as in Fig. 3. Blue circles show 50 replicate simulations, and red crosses show expected values.

**Figure 7:**
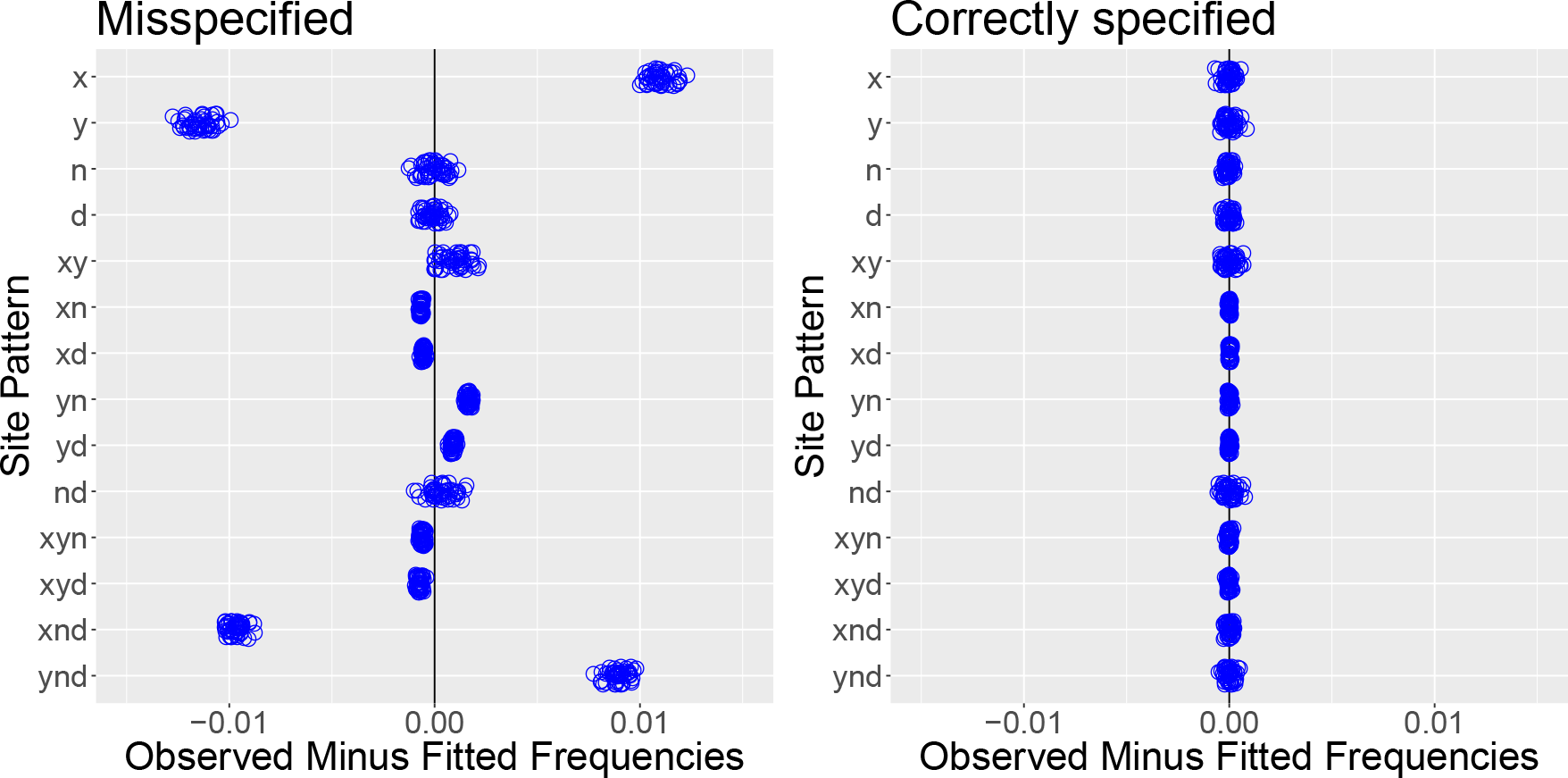
Residuals from misspecified and correctly-specified models. Each circle represents one of the simulated data sets in Fig. 3. The misspecified model ignores the two episodes of gene flow seen in Fig. 1.

Table 1 compares the two models and shows that the one with *N* → *Y* gene flow is unambiguously better than the one without gene flow. However, the residuals of this new model (not shown) still show discrepancies, which might lead us to consider adding *D* → *Y* gene flow to the model. Table 2 shows that this third model is unambiguously better than the one with only one episode of gene flow. The residuals (right panel of Fig. 7) show that this model provides a good description of the data. In this example, the correct model was identifiable because the alternate models could not fully account for the pattern in the data.

**Table 1:** Booma weights for models with and without *N* → *Y* gene flow. All models re-express free variables in terms of principal components. Models with reduced dimension exclude principal components that explain less than a fraction 0.001 of the variance.

| Weights |  | Model |
| --- | --- | --- |
| bepe | clic |  |
| 0 | 0 | No gene flow; full dimension |
| 0 | 0 | No gene flow; reduced dimension |
| 0.04 | 0.5 | $N \rightarrow Y$ gene flow; full dimension |
| 0.96 | 0.5 | $N \rightarrow Y$ gene flow; reduced dimension |

**Table 2:** Booma weights for models with and without *D* → *Y* gene flow. All models include *N* → *Y* gene flow and re-express free variables in terms of principal components. Models with reduced dimension exclude principal components that explain less than a fraction 0.001 of the variance.

| Weights |  | Model |
| --- | --- | --- |
| bepe | clic |  |
| 0 | 0 | No $D \rightarrow Y$ gene flow; full dimension |
| 0 | 0 | No $D \rightarrow Y$ gene flow; reduced dimension |
| 0.42 | 0.12 | $D \rightarrow Y$ gene flow; full dimension |
| 0.58 | 0.88 | $D \rightarrow Y$ gene flow; reduced dimension |

There are also less tractable identifiability problems. Let us consider two. Figure 8 shows a model that is like that in the simulations (Fig. 1) but has an additional episode of gene flow from a “superarchaic” population (*S*) into Denisovans (*D*), as suggested by Prüfer et al Prüfer et al. [2014]. When the superarchaic admixture fraction is zero, this model reduces to that used in our simulations. As expected, legofit’s estimate of this parameter was very close to zero in all simulation replicates, and all other parameters were also well estimated. Consequently, this model provides an excellent fit to the data, comparable to that in the right panel of Fig. 7. Nonetheless, I expected bepe and clic to prefer the correct model because of its simplicity. Instead, bepe and clic gave appreciable weight to both models but preferred the more complex one, as shown in table 3. This did not lead to incorrect inferences, because all parameters were well estimated.

**Table 3:** Booma weights for models with and without superarchaic admixture. All models include *N* → *Y* and *D* → *Y* gene flow and re-express free variables in terms of principal components. Models with reduced dimension exclude principal components that explain less than a fraction 0.001 of the variance.

| Weights |  | Model |
| --- | --- | --- |
| bepe | clic |  |
| 0.24 | 0.04 | No superarchaic admixture; full dimension |
| 0.02 | 0.16 | No superarchaic admixture; reduced dimension |
| 0 | 0 | Superarchaic admixture; full dimension |
| 0.74 | 0.80 | Superarchaic admixture; reduced dimension |

**Figure 8:**
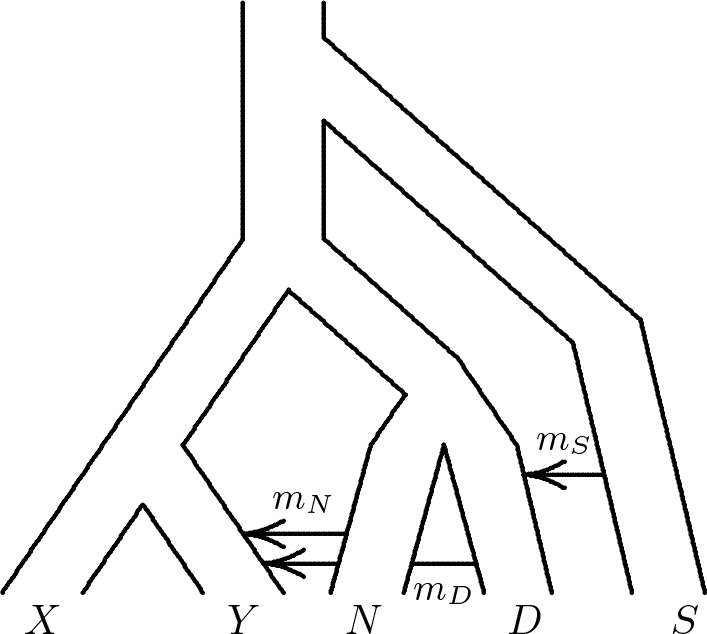
Admixture from a superarchaic population (*S*) into Denisovans (*D*).

Table 4 illustrates another identifiability problem. It compares the standard model (Fig. 1) with one in which the order of the two admixture events is reversed: *D* → *Y* admixture precedes *N* → *Y* admixture. This change has little effect on site pattern frequencies, and all parameters are well estimated. I expected bepe and clic to weight these models roughly equally. The table shows that they do give appreciable weight to both models but prefer the (incorrect) reversed model. In another experiment (not shown), using ms instead of msprime, bepe gave 94% of the weight to the true model. Bepe and clic both behave sensibly when dealing with models that are indistinguishable or nearly so. In such cases, they tend to give appreciable weight to several models. We cannot assume, however, that they will always prefer the correct model.

**Table 4:**
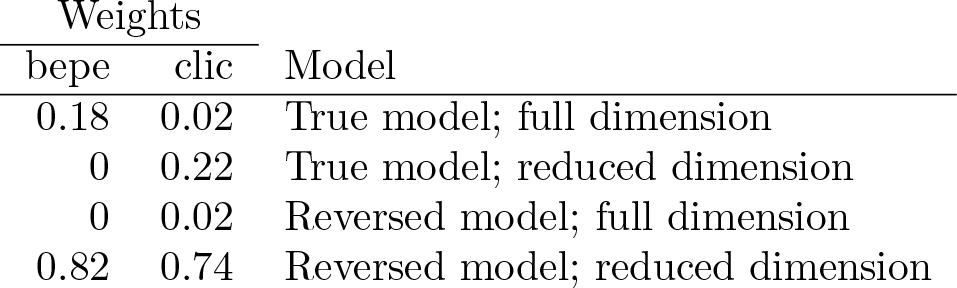
Booma weights for models with and without reversing the order of the two admixture events in Fig. 1. All models include *N* → *Y* and *D* → *Y* gene flow and re-express free variables in terms of principal components. Models with reduced dimension exclude principal components that explain less than a fraction 0.001 of the variance.

## Discussion

There are two reasons for studying site patterns rather than the full site frequency spectrum, the first of which involves statistical power at deep time scales. As we look backwards into the past, large samples coalesce rapidly to small collections of ancestors. For this reason, although large samples are essential for recent history, their value is limited in the distant past. Furthermore, the random-haploid samples used by legofit provide an advantage: they insulate the analysis from recent population history. If we had sampled several haploid genomes from population *X* in Fig. 1, then our model would need parameters describing changes in the size of *X* since its separation from *Y*. With legofit, these parameters aren’t needed, because no coalescent events can occur until *X* and *Y* merge into their ancestral population. Thus, site pattern frequencies reduce the parameter count without losing much power at deep time scales. They are most valuable for studying the deep history of multiple populations.

## Conclusions

The Legofit package provides computer programs for estimating population histories. It uses the frequencies of nucleotide site patterns to summarize genetic data. The package includes programs that tabulate these frequencies, calculate their expected values, and use them to estimate parameters describing population history. It includes facilities for model selection and model averaging. It uses principal components to reduce the complexity of high-dimensional models of history. Legofit outperforms several methods that have been widely used to study archaic admixture in humans.

## Availability and requirements

|  |  |
| --- | --- |
| Project name | Legofit |
| Home page | https://github.com/alanrogers/legofit |
| OS | Linux and macOS |
| Language | C and Python |
| License | Internet Systems Consortium License |
| Requirements | pthread and the Gnu Scientific Library |
| Data | available at datadryad.org |

## Acknowledgements

I am grateful to Alan Achenbach, Kiela Gwin, Nathan Harris, Louise Holbrook, Mitchell Lokey, and Daniel Tabin, who have all used the software and provided feedback. Daniel Tabin helped write several programs within the package. Elizabeth Cashdan, Ilan Gronau, Timothy Webster provided useful comments on the text. The package makes use of tinyexpr, which was written by Lewis Van Winkle. Legofit’s implementation of the differential evolution algorithm is based on that of Rainer Storn and Ken Price.

## Funding

This work was supported by NSF award BCS 1638840 and by the Center for High Performance Computing at the University of Utah.

