## Supplementary Materials for "Legofit: Estimating Population History from Genetic Data"

### Additional File
### Legofit: Estimating Population History from Genetic Data

Alan R. Rogers*

February 11, 2019

#### 1 Model

This additional file documents the analyses discussed in the main text. Because few readers will be familiar with Legofit, I describe these analyses in detail and provide selected input and output files as appendices. All input and output files are archived at `datadryad.org`. Within this archive, analyses are organized in several directories, each of which has its own appendix in this document. The directories and corresponding appendices are "legosim" (appendix A), "msp" (for msprime analyses, appendix B), "ms" (for ms analyses, appendix C), "scrm" (for scrm analyses, appendix D), and "treemix" (for TreeMix analyses, appendix E).

The model, illustrated in Fig. 1 of the main text, describes the history of four haploid samples, one each from populations $X$, $Y$, $N$, and $D$, which are meant to represent an African population, a Eurasian population, Neanderthals, and Denisovans. The population ancestral to $X$ and $Y$ is called $XY$, that ancestral to $N$ and $D$ is $ND$, and that ancestral to all four populations is $XYND$. These populations differ in size, and I use notation such as $2N_{XY}$ to represent their sizes. There is gene flow from $N \to Y$ at time $T_{m_N}$ and from $D \to Y$ at time $T_{m_D}$.

The parameters of this model are listed in table 1. The parameters fall into three categories: fixed, free, and constrained. Fixed parameters are constants. Free parameters are estimated by legofit but behave as constants during computer simulations. Constrained parameters are functions of one or more other parameters and are not estimated. One separation time, $T_{XYND}$, is treated as a fixed parameter in order to calibrate the molecular clock. The sizes of populations $X$ and $Y$ do not affect site pattern frequencies, because no coalescent events can occur within them. Consequently, $2N_X$ and $2N_Y$ are defined as fixed constants. Parameters $T_{m_N}$ and $T_{m_D}$ have little effect on site pattern frequencies, so they are also treated as constants.

The .lgo file defining this model is called `true.lgo` and is listed below in appendix A.1 on p. 6. The syntax of this file is described in the Legofit documentation. I used the following command to estimate expected values under this model:

```
legosim -1 -i 100000000 true.lgo > true.patfrq
```

Here, `-1` tells legosim to include singleton site patterns, and `-i` sets to $10^8$ the number of iterations to use in estimating expected values. The output is in appendix A.2 on p. 7.

*Dept. of Anthropology, 260 South Central Campus Dr., Rm. 4428, University of Utah, Salt Lake City, UT 84112.

Table 1: Parameters of model

| | | |
|---|---|---|
| *Time parameters (generations ago)* | | |
| $T_{XYND} = 25920$ | fixed | separation time of $XY$ and $ND$ |
| $T_{XY} = 3788$ | free | separation time of $X$ and $Y$ |
| $T_{ND} = 15000$ | free | separation time of $N$ and $D$ |
| $T_N = 1760$ | free | age of Neanderthal fossil |
| $T_D = 1734$ | free | age of Denisovan fossil |
| $T_{m_N} = 1897$ | fixed | time of $N \to Y$ gene flow |
| $T_{m_D} = T_{m_N} - 1$ | constrained | time of $D \to Y$ gene flow |
| *Haploid population sizes* | | |
| $2N_{XYND} = 64964.1$ | free | size of population $XYND$ |
| $2N_{XY} = 44869.2$ | free | size of population $XY$ |
| $2N_{ND} = 5000$ | free | size of population $ND$ |
| $2N_X = 20000$ | fixed | size of population $X$ |
| $2N_Y = 20000$ | fixed | size of population $Y$ |
| $2N_N = 9756.8$ | free | size of population $N$ |
| $2N_D = 5000$ | free | size of population $D$ |
| *Admixture fractions* | | |
| $m_N = 0.05$ | free | fraction of $Y$ replaced by immigrants from $N$ |
| $m_D = 0.025$ | free | fraction of $Y$ replaced by immigrants from $D$ |

#### 2 Simulations

I generated 50 simulated data sets using each of three simulation programs: msprime [2], ms [1], and scrm [6]. In all three cases, the simulations used the same model of history, with the same parameter values, as described above.

I ran msprime within a Python script, which is listed in appendix B.1 on p. 8. The Python script was launched by a shell script, sim.sh (appendix B.2 on p. 11), which was executed as follows:

```
seq 0 49 | xargs -n 1 -P 16 bash sim.sh
```

In this command, seq, xargs, and bash are Unix/Linux/macOS commands. The command generates 50 output files with names like `sim0.opf`, `sim1.opf`, and so on. One of these is shown in appendix B.3 on p. 11.

ms is a command-line program, which I execute under the control of a Python script, ms.py (appendix C.1 on p. 28). For each simulation replicate, ms.py is launched by a bash script, sim.sh (appendix C.2 on p. 31). This script was launched for each of 50 simulation replicates as follows:

```
seq 0 49 | xargs -n 1 -P 16 bash sim.sh
```

scrm is also a command-line program. As with ms, I execute it under control of a Python script, scrm.py (appendix D.1 on p. 31). Rather than doing this on my own workstation, I did it on a compute cluster at the University of Utah's Center for High Performance Computing. This involves using a slurm script (appendix D.2 on p. 33), which is executed on the CHPC server as follows:

```
sbatch --array=0-49 sim.slr
```

This queues 50 jobs, one for each simulation replicate.

### 3 Graphing site pattern frequencies

To make graphs such as Fig. 6 of the main text, I used an R script. The script from the msprime directory is in appendix B.4 on p. 12. It was executed as follows:

```
Rscript patfrq.r
```

This produces two files, msp-dotfrq.pdf and msp-doterr.pdf. Fig. 2 of the main text shows the second of these, along with the corresponding output of ms and scrm. These plots show that all three simulators generate site pattern frequencies that agree with the expectations calculated by legosim. Furthermore, they show that the three simulators generate similar sampling distributions.

### 4 Estimation

I will describe the procedure used to estimate parameters from msprime simulations. The same procedure was used with ms and scrm.

#### 4.1 Begin with an unconstrained analysis

The first step is to create a.lgo (appendix B.5, p. 13), which is identical to true.lgo (described above in sec. 1) except that the free parameters are arbitrarily set to incorrect values, so that legofit will have work to do. The first stage of analysis on this file uses a slurm script called a1.slr (appendix B.6, p. 14), which is executed on the CHPC cluster as follows:

```
sbatch --array=0-49 a1.slr
```

This generates 50 files with names like a1-0.legofit and 50 with names like a1-0.state, where "0" is the index of the simulation replicate. The .legofit files describe the estimates obtained from the simulated data sets, and the .state files record the state of the differential evolution algorithm at the end of the run. The .state files make it possible to restart the estimation procedure where it left off.

The second stage in the estimation deals with the possibility that the composite likelihood function may have multiple local optima. If it does, the various .legofit files may be at different local optima. Stage 2 of the analysis allows legofit to choose among these. It does so by using legofit's `--stateIn` option to read in all 50 of the .state files produced during stage 1. This allows legosim to construct an initial swarm of points that is distributed across all of the solutions found during stage 1. The slurm script for stage 2 is a2.slr (appendix B.7, p. 15). Note that the `--stateIn` option is used many times. This slurm script is launched as follows:

```
sbatch --array=0-49 a2.slr
```

This produces 50 output files with names like a2-0.legofit. No .state files are produced, because a2.slr does not use the `--stateOut` option.

Next, I use bepe to calculate the bootstrap estimate of predictive error:

```
export LC_ALL=C
bepe sim*.opf -L a2-*.legofit > a2.bepe
```

The first line is a bash command that makes sure a consistent sort order is used. Otherwise, when `sim*.opf` and `a2-*.legofit` are expanded into lists of file names, they will appear in an order that depends on the locale setting of the operating system. This will affect the order of lines in the output .bepe file. For this reason, .bepe files may be inconsistent if they are produced on machines with different locale settings. The `LC_ALL=C` setting ensures consistency. The output, a2.bepe (appendix B.8, p. 16) contains a bepe value for each simulation replicate.

The procedure for clic is similar:

```
export LC_ALL=C
clic a-sim*-2.pts -L a2-*.legofit > a2.clic
```

The resulting file is a2.clic (appendix B.9, p. 17).

The following command generates a flat file with a row for each simulation replicate and a column for each parameter:

```
export LC_ALL=C
flatfile.py a2-*.legofit > a2.flat
```

These flat files are needed by booma. They can also be useful when importing the data into other packages to do further statistical analysis or make graphs.

Fig. 3 of the main text is a scatter-plot matrix, with a sub-plot for each pair of parameters. It was made with the R script a2.r (appendix B.10, p. 19).

##### 4.2 Using principal components to reduce dimensionality

Having finished an unconstrained analysis of these data, we now re-express the free variables in terms of principal components (PCs). This makes legofit's job easier, because it is working with uncorrelated variables. We can also reduce the number of dimensions.

Here is a command that generates a new .lgo file, which re-expresses all the free parameters as linear combinations of PCs:

```
(grep ^# a.lgo; pclgo a.lgo a2-*.legofit; grep -v ^# a.lgo | \
egrep -v "\<free\>") > b.lgo
```

The first grep command in this pipeline extracts all the comments from `a.lgo`. They will end up at the top of the new .lgo file. Then, pclgo generates output that defines new free variables called pc1, pc2, and so on, and re-expresses all the free variables of `a.lgo` in terms of these new variables. The rest of the pipeline prints all the lines in `a.lgo` *except* comments and definitions of free variables. These lines go at the end of the new .lgo file. The resulting file is b.lgo (appendix B.11, p. 19). It expresses the 11 free parameters in terms of all 11 PCs, so there is no reduction of dimensions.

To reduce the number of dimensions, use the `--tol` argument of pclgo:

```
(grep ^# a.lgo; pclgo --tol 0.001 a.lgo a2-*.legofit; \
grep -v ^# a.lgo | egrep -v "\<free\>") > c.lgo
```

This says to include only those principal components that explain at least a fraction 0.001 of the variance. There are 9 PCs that satisfy this criterion, so c.lgo (appendix B.12, p. 21) expresses the 11 free variables in terms of 9 principal components.

The analysis of b.lgo and c.lgo is exactly like that described above for a.lgo. There is a first stage in which the `--stateOut` option of legofit is used to generate a .state file for each simulation replicate. Then in the second stage of analysis, multiple `--stateIn` options are used. In the end, we get legofit output files from stage 2 with names like b2-0.legofit and c2-0.legofit. These are used with bepe and flatfile.py to generate b2.bepe, c2.bepe, b2.flat, and c2.flat.

The last stage in the analysis uses booma to do bootstrap model averaging:

```
booma a2.bepe b2.bepe c2.bepe -F a2.flat b2.flat c2.flat > a-b-c.bma
```

The output file is msp/a-b-c.bma. Near the top of this file, you will find the lines:

```
#         Weight        MSC_file       Flat_file
#              0         a2.bepe         a2.flat
#           0.42         b2.bepe         b2.flat
#           0.58         c2.bepe         c2.flat
```

This says that the unconstrained analysis (based on a.lgo) got weight 0 and can therefore be ignored. The other two analyses (both using PCs to re-express variables) got positive weights. The rest of the .bma file contains model-averaged estimates of the free parameters.

To graph these results, use the R code in bma.r (appendix B.13, p. 23):

```
Rscript bma.r
```

This produces several output files. Of these, `msp-bma-mdot.pdf`, `msp-bma-ndot.pdf`, and `msp-bma-tdot.pdf` are used in Fig. 4 of the main text.

### 5 Residual plots

Figure 7 of the main text has residual plots for two models, one misspecified and the other correctly specified. Each of these was based on an analysis like that described in section 4, so I will present here only the R code that made the residual plot for the misspecified model. This code is in nma2resid.r (appendix B.14, p. 26).

### 6 Previously-published estimators

The TreeMix analysis uses simulated data generated by treemix.py (appendix E.2, p. 34). This uses msprime to do simulations identical to those discussed above and in appendix B.1, but writes the output in TreeMix format. The steps in this analysis are detailed in README.md (appendix E.1, p. 34) and treemix.sh (appendix E.3, p. 38).

Statistics "Nea" and "Den" were defined by Patterson et al. [4] to estimate (in their notation) $f_1 - f_2$, where $f_1$ is the fraction of archaic admixture into population $H_1$ and $f_2$ is that into $H_2$. In my notation, $H_1$ becomes $Y$, $H_2$ becomes $X$. I also assume that $f_2 = 0$. Nea and Den estimate admixture from Neanderthals and Denisovans, respectively.

Nea and Den are defined [4, Eqns. 5 and 8, p. 42] as

$$\begin{aligned}
\text{Nea}(H_1, H_2) &= \frac{\sum_{i=1}^{N} [p^i_{H_1}(1 - p^i_{H_2}) - (1 - p^2_{H_1})p^i_{H_2}]p^i_{\text{Denisova}}}{\sum_{i=1}^{N} [p^i_{\text{Neandertal}}(1 - p^i_{H_2}) - (1 - p^i_{\text{Neandertal}})p^i_{H_2}]p^i_{\text{Denisova}}} \\
\text{Den}(H_1, H_2) &= \frac{\sum_{i=1}^{N} [p^i_{H_1}(1 - p^i_{H_2}) - (1 - p^i_{H_1})p^i_{H_2}]p^i_{\text{Neandertal}}}{\sum_{i=1}^{N} [p^i_{\text{Denisova}}(1 - p^i_{H_2}) - (1 - p^i_{\text{Denisova}})p^i_{H_2}]p^i_{\text{Neandertal}}}
\end{aligned}$$

The allele frequencies are defined [4, p. 37] such that $p^i_K$ is the fraction of reads from sample $K$ that carry the derived allele at site $i$.

To re-express these definitions in terms of nucleotide site patterns, write $X$ for $H_2$, $Y$ for $H_1$, $N$ for Neandertal, $D$ for Denisovan, and $E[]$ for the operation of averaging across sites. Let $p_k = 1 - q_k$

represent the fraction of derived alleles in the sample of some nucleotide site from population $k$. Then the formulas above become

$$\begin{aligned}\text{Nea}(Y,X) &= \frac{E[q_X p_Y p_D - p_X q_Y p_D]}{E[q_X p_N p_D - p_X q_N p_D]}\\ \text{Den}(Y,X) &= \frac{E[q_X p_Y p_N - p_X q_Y p_N]}{E[q_X p_N p_D - p_X p_N q_D]}\end{aligned}$$

Let $P_m$ denote the relative frequency of site pattern $m$ in a tabulation of all four samples. Then

$$\begin{aligned}\text{Nea}(Y,X) &= \frac{P_{yd} + P_{ynd} - P_{xd} - P_{xnd}}{P_{nd} + P_{ynd} - P_{xd} - P_{xyd}}\\ \text{Den}(Y,X) &= \frac{P_{yn} + P_{ynd} - P_{xn} - P_{xnd}}{P_{nd} + P_{ynd} - P_{xn} - P_{xyn}}\end{aligned}$$

These formulas are implemented in neaden.py (appendix B.15, p. 26).

The Nea and Den values for each simulation replicate were calculated as follows:

```
./neaden.py sim*.opf > neaden.out
```

This same Python script was also used to calculate the expected values of Nea and Den:

```
./neaden.py ../legosim/true.patfrq
```

The result agrees with the theory of Rogers and Bohlender [5]. This calculation approximates the expection of a ratio by the corresponding ratio of expected values. With genome-scale data, the approximation is excellent, because the law of large numbers ensures that the numerator and denominators will both be close to their expectations. Fig. 5 of the main text was generated using neaden.r (appendix B.16, p. 28):

```
Rscript neaden.r
```

This command generates neaden.pdf.

### A legosim

#### A.1 true.lgo

```
# Model: ((X,Y), (N,D)), migration: N -> Y, D -> Y
time fixed   zero  = 0
time fixed   Txynd = 25920   # \citet[table~S12.2, p.~90]{Li:N-505-43-S88}
time free      Tnd = 15000
time free      Txy = 3788    # Schiffels and Durbin fig 4
time fixed     TmN = 1897    # \citep[table~2]{Sankararaman:PLO-8-e1002947}
time free      Td  = 1734    # fossil \citep[p.~43]{Prufer:N-505-43}
time free      Ta  = 1760    # altai fossil: slightly older
twoN free   twoNnd = 5000
twoN free    twoNn = 9756.8  # Fig~1,left,Rogers:PNA-114-E10258
twoN free    twoNd = 5000
twoN free   twoNxy = 44869.2 # Fig~1,left,Rogers:PNA-114-E10258
```

```
twoN fixed   twoNx = 20000
twoN fixed   twoNy = 20000
twoN free twoNxynd = 64964.1 # Fig~1,left,Rogers:PNA-114-E10258
mixFrac free    mN = 0.05    # arbitrary
mixFrac free    mD = 0.025   # arbitrary
time constrained TmD = Td - 1 # arbitrary
segment x    t=zero   twoN=twoNx   samples=1
segment y    t=zero   twoN=twoNy   samples=1
segment n    t=Ta     twoN=twoNn   samples=1
segment n2   t=TmN    twoN=twoNn
segment d0   t=TmD    twoN=twoNd
segment d    t=Td     twoN=twoNd   samples=1
segment y1   t=TmD    twoN=twoNy
segment y2   t=TmN    twoN=twoNy
segment nd   t=Tnd    twoN=twoNnd
segment xy   t=Txy    twoN=twoNxy
segment xynd t=Txynd  twoN=twoNxynd
mix    y  from y1 + mD * d0
mix    y1 from y2 + mN * n2
derive x  from xy
derive y2 from xy
derive n  from n2
derive n2 from nd
derive d0 from d
derive d  from nd
derive xy from xynd
derive nd from xynd
```

##### A.2 true.patfrq

```
############################################################
# legosim: generate site patterns by coalescent simulation #
#                       version 1.67                       #
############################################################

# Program was compiled: Nov 18 2018 15:29:32
# Program was run: Fri Nov 23 10:58:57 2018

# cmd: legosim -1 -i 100000000 true.lgo
# nreps                       : 100000000
# input file                  : true.lgo
# not simulating mutations
# including singleton site patterns.
#       SitePat E[BranchLength]
              x   49997.8953109
              y   44176.5940544
              n   21184.8632778
              d   21403.3689990
```

```
     x:y   38446.7112987
     x:n     763.9517961
     x:d     793.4660017
     y:n    1366.3376042
     y:d    1176.5437076
     n:d   50156.2573985
   x:y:n    1674.0450624
   x:y:d    1641.8190321
   x:n:d   11576.5956204
   y:n:d   16412.4333629
```

### B Msprime

#### B.1 msp.py

```
#!/usr/bin/python3
import msprime
import os, sys, time

def usage():
    print("Usage: ./msp.py [options]")
    print("  where options may include:")
    print("  -r or --run: run simulation. Default: run")
    print("               DemographyDebugger")
    sys.exit(1)

do_simulation = False
for arg in sys.argv[1:]:
    if arg == "-r" or arg == "--run":
        do_simulation = True
    else:
        usage()

# time parameters in generations
Txynd = 25920
Tnd = 15000
Txy = 3788
Td = 1734      # age of Denisova fossil
Ta = 1760      # age of Altai fossil
TmN = 1897     # time of Neanderthal admixture
TmD = Td-1     # time of Denisovan admixture

# haploid population sizes
Nxynd = 64964.1/2.0  # ancestral population
Nxy = 44869.2/2.0
Nnd = 5000/2.0
Nn = 9756.8/2.0
Nd = 5000/2.0
```

```
Nx = 20000/2.0 # modern Africa
Ny = 20000/2.0 # modern Europe

# admixture
mN = 0.05
mD = 0.025

nchromosomes = 1000      # number of chromosomes
basepairs = 2e6   # number of nucleotides per chromosome
u_per_site = 1.4e-8 # mutation

# Recombination rate.  recomb is the probability of recombination
# between sites at opposite ends of the simulated sequence.
c = 1e-8 # rate per base pair per generation

# One haploid sample from each of 4 populations: two modern (X,Y),
# and two archaic (N,D).
samples = [
    msprime.Sample(population=0, time=0),  # population X
    msprime.Sample(population=1, time=0),  # population Y
    msprime.Sample(population=2, time=Ta), # population N
    msprime.Sample(population=3, time=Td)  # population D
]

lbl = ("x", "y", "n", "d")
npops = len(lbl)

# associate sample sizes with population labels
sampsize = {}
for s in lbl:
    sampsize[s] = 0

for samp in samples:
    ndx = samp.population
    assert ndx < len(lbl)
    assert lbl[ndx] in sampsize
    sampsize[lbl[ndx]] = 1

for s in lbl:
    if not (sampsize[s] > 0):
        print("Population %s has no samples" % s, file=sys.stderr)
        sys.exit(1)

# Population configurations. No sample sizes are listed here, because
# those are specified above in "samples".
popconf = [
    msprime.PopulationConfiguration(initial_size=Nx),
    msprime.PopulationConfiguration(initial_size=Ny),
```

```
    msprime.PopulationConfiguration(initial_size=Nn),
    msprime.PopulationConfiguration(initial_size=Nd)
]

events = [
    msprime.MassMigration(
        time=TmD,
        source=1,
        dest=3,
        proportion=mD), # D->Y gene flow
    msprime.MassMigration(
        time=TmN,
        source=1,
        dest=2,
        proportion=mN), # N->Y gene flow
    msprime.MassMigration(
        time=Txy,
        source=1,
        dest=0,
        proportion=1.0), # X-Y split
    msprime.PopulationParametersChange(
        time=Txy,
        initial_size=Nxy,
        population_id=0),
    msprime.MassMigration(
        time=Tnd,
        source=3,
        dest=2,
        proportion=1.0), # N-D split
    msprime.PopulationParametersChange(
        time=Tnd,
        initial_size=Nnd,
        population_id=2),
    msprime.MassMigration(
        time=Txynd,
        source=2,
        dest=0,
        proportion=1.0), # XY-ND split
    msprime.PopulationParametersChange(
        time=Txynd,
        initial_size=Nxynd,
        population_id=0)
]

if do_simulation:
    # run simulation
    seed = int(time.time()) ^ os.getpid()
    sim = msprime.simulate(samples = samples,
```

```
                            population_configurations = popconf,
                            demographic_events = events,
                            length = basepairs,
                            recombination_rate = c,
                            mutation_rate = u_per_site,
                            num_replicates = nchromosomes,
                            random_seed = seed)

    # header
    print("npops = %d" % len(lbl))
    print("%s %s" % ("pop", "sampsize"))
    for s in lbl:
        print("%s %d" % (s, sampsize[s]))

    for i, chromosome in enumerate(sim):
        for variant in chromosome.variants():
            print(i, end=" ")
            for g in variant.genotypes:
                print(g, end=" ")
            print()

else:
    # Run demography debugger and quit
    dd = msprime.DemographyDebugger(
        population_configurations=popconf,
        demographic_events=events)
    dd.print_history()
    print("Use \"./sim -r\" to run simulation")
```

##### B.2 sim.sh

```
# msp.py is a Python script that executes msprime and generates output in the "sim"
# format, which is described in the document of simpat, within the Legofit package.
# simpat is a program that reads sim format and tabulates site patterns.
ofile=sim${1}.opf
efile=sim${1}.err
python3 msp.py -r | simpat 1>${ofile} 2>${efile}
```

##### B.3 sim0.opf

```
# simpat version 1.67
# Including singleton site patterns.
# Number of site patterns: 14
# Nucleotide sites: 7298790
# Sites used: 7298790
#      SitePat             E[count]
              x      1401902.0000000
```

```
            y      1239084.0000000
            n       589582.0000000
            d       598305.0000000
          x:y      1074236.0000000
          x:n        21136.0000000
          x:d        21693.0000000
          y:n        38330.0000000
          y:d        31022.0000000
          n:d      1408276.0000000
        x:y:n        45407.0000000
        x:y:d        44519.0000000
        x:n:d       324614.0000000
        y:n:d       460684.0000000
```

#### B.4 patfrq.r

```
library(ggplot2)

# Adjust text size. Default is 11
mytheme = theme_get()
mytheme$text$size = 25
theme_set(mytheme)

# Construct a script, which will be processed by the linux command
# "sed". The script has two commands separated by a semicolon. The
# first command (s/[#:]//g) deletes sharps and colons from the
# input. The second command (beginning with /SitePat/) tells sed to
# print lines from the one containing "SitePat" to the end of the
# file.
script <- "'s/[#:]//g; /SitePat/,$ p'"

# Create a data frame with 0 rows and 2 columns, with labels "pat" and "frq"
df <- data.frame(pat=vector("character", 0), frq=vector("numeric",0))

# read data files and concatenate into df
for(i in 0:49) {
    # file name
    fname <- paste("sim", i, ".opf", sep="")

    # linux shell command.
    cmd <- paste("sed -n", script, fname)

    # Open a pipe that executes the shell command. read.table reads from
    # this pipe as though it were a file. Then close the pipe.
    p <- pipe(cmd, open="r")
    df2 <- read.table(p, header=T)
    close(p)
```

```
    # column names
    names(df2) <- c("pat", "frq")

    # make "pat" an order vector so that the site patterns will
    # plot in the same order as the input file.
    df2$pat <- ordered(df2$pat, levels=rev(df2$pat))

    # convert to relative frequencies
    df2$frq <- df2$frq / sum(df2$frq)

    # append to df
    df <- rbind(df, df2)
}

# Data frame "tru" contains true parameter values
cmd <- paste("sed -n", script, "../legosim/true.patfrq")
p <- pipe(cmd, open="r")
tru <- read.table(p, header=T)
close(p)
names(tru) <- c("pat", "true")
tru$pat <- ordered(tru$pat, levels=rev(tru$pat))
tru$true <- tru$true / sum(tru$true)

df2 <- merge(df, tru)
ggplot(df2, aes(frq, pat)) +
    ggtitle("msprime") +
    xlab("Frequency") + ylab("Site Pattern") +
    geom_jitter(height=0.2, width=0, shape=1, alpha=0.333, color="blue",
                size=3) +
    geom_point(mapping=aes(true, pat), shape=4, color="red",
               size=3)
ggsave("msp-dotfrq.pdf")

df2$err <- df2$frq - df2$true
ggplot(df2, aes(err, pat)) +
    ggtitle("msprime") +
    xlab("Error") + ylab("Site Pattern") +
    geom_vline(xintercept=0) +
    xlim(c(-0.0016, 0.0016)) +
    geom_jitter(height=0.2, width=0, shape=1, color="blue",
                size=3)
ggsave("msp-doterr.pdf")
```

##### B.5 a.lgo

```
# Model: ((X,Y), (N,D)), migration: N -> Y, D -> Y
# This file is like true.lgo but the free parameter values have been
# arbitrarily moved to incorrect values so that legofit will have some
```

```
# work to do.
time fixed   zero  = 0
twoN fixed   one   = 1
time fixed   Txynd = 25920   # \citet[table~S12.2, p.~90]{Li:N-505-43-S88}
time free      Tnd = 20000
time free      Txy = 2500
time fixed     TmN = 1897    # \citep[table~2]{Sankararaman:PLO-8-e1002947}
time fixed     TmD = 1       # arbitrary
time free      Td  = 1200
time free      Ta  = 1100
twoN free   twoNnd = 800
twoN free    twoNn = 1000
twoN free    twoNd = 1000
twoN free   twoNxy = 20000
twoN free twoNxynd = 20000
mixFrac free    mN = 0.01
mixFrac free    mD = 0.01
segment x    t=zero   twoN=one     samples=1
segment y    t=zero   twoN=one     samples=1
segment n    t=Ta     twoN=twoNn   samples=1
segment n2   t=TmN    twoN=twoNn
segment d0   t=TmD    twoN=one
segment d    t=Td     twoN=twoNd   samples=1
segment y1   t=TmD    twoN=one
segment y2   t=TmN    twoN=one
segment nd   t=Tnd    twoN=twoNnd
segment xy   t=Txy    twoN=twoNxy
segment xynd t=Txynd  twoN=twoNxynd
mix    y  from y1 + mD * d0
mix    y1 from y2 + mN * n2
derive x  from xy
derive y2 from xy
derive n  from n2
derive n2 from nd
derive d0 from d
derive d  from nd
derive xy from xynd
derive nd from xynd
```

##### B.6 a1.slr

```
#!/bin/bash
#SBATCH -J a1
#SBATCH --account=rogersa-kp
#SBATCH --partition=rogersa-kp
#SBATCH --time=36:00:00
#SBATCH --nodes 1
#SBATCH --ntasks 1
```

```
#SBATCH -o a1-%a.legofit # stdout
#SBATCH -e a1-%a.err # stderr

i=${SLURM_ARRAY_TASK_ID}
ifile=‘printf "sim%d.opf" $i‘    # input file
stateout=‘printf "a1-%d.state" $i‘

time legofit -1 --stateOut ${stateout} --tol 3e-5 \
  -S 5000@10000 -S 100@100000 -S 1000@2000000 a.lgo ${ifile}
```

##### B.7 a2.slr

```
#!/bin/bash
#SBATCH -J a2
#SBATCH --account=rogersa-kp
#SBATCH --partition=rogersa-kp
#SBATCH --time=36:00:00
#SBATCH --nodes 1
#SBATCH --ntasks 1
#SBATCH -o a2-%a.legofit # stdout
#SBATCH -e a2-%a.err # stderr

i=${SLURM_ARRAY_TASK_ID}
ifile=‘printf "sim%d.opf" $i‘    # input file
time legofit -1 --tol 2e-5 \
  -S 1000@2000000 a.lgo ${ifile} \
  --stateIn a1-0.state \
  --stateIn a1-1.state \
  --stateIn a1-10.state \
  --stateIn a1-11.state \
  --stateIn a1-12.state \
  --stateIn a1-13.state \
  --stateIn a1-14.state \
  --stateIn a1-15.state \
  --stateIn a1-16.state \
  --stateIn a1-17.state \
  --stateIn a1-18.state \
  --stateIn a1-19.state \
  --stateIn a1-2.state \
  --stateIn a1-20.state \
  --stateIn a1-21.state \
  --stateIn a1-22.state \
  --stateIn a1-23.state \
  --stateIn a1-24.state \
  --stateIn a1-25.state \
  --stateIn a1-26.state \
  --stateIn a1-27.state \
  --stateIn a1-28.state \
```

```
  --stateIn a1-29.state \
  --stateIn a1-3.state \
  --stateIn a1-30.state \
  --stateIn a1-31.state \
  --stateIn a1-32.state \
  --stateIn a1-33.state \
  --stateIn a1-34.state \
  --stateIn a1-35.state \
  --stateIn a1-36.state \
  --stateIn a1-37.state \
  --stateIn a1-38.state \
  --stateIn a1-39.state \
  --stateIn a1-4.state \
  --stateIn a1-40.state \
  --stateIn a1-41.state \
  --stateIn a1-42.state \
  --stateIn a1-43.state \
  --stateIn a1-44.state \
  --stateIn a1-45.state \
  --stateIn a1-46.state \
  --stateIn a1-47.state \
  --stateIn a1-48.state \
  --stateIn a1-49.state \
  --stateIn a1-5.state \
  --stateIn a1-6.state \
  --stateIn a1-7.state \
  --stateIn a1-8.state \
  --stateIn a1-9.state
```

##### B.8 a2.bepe

```
################################################
# bepe: bootstrap estimate of predictive error #
#                version 1.82                  #
################################################

# Program was compiled: Feb 10 2019 10:16:21
# Program was run: Sun Feb 10 10:24:34 2019
# Correcting for bootstrap bias

#          bepe        DataFile     LegofitFile
2.226031606e-07        sim0.opf    a2-0.legofit
2.816382566e-07        sim1.opf    a2-1.legofit
2.945104273e-07       sim10.opf   a2-10.legofit
2.025672297e-07       sim11.opf   a2-11.legofit
2.236214047e-07       sim12.opf   a2-12.legofit
3.275322083e-07       sim13.opf   a2-13.legofit
1.738806689e-07       sim14.opf   a2-14.legofit
```

```
2.222293762e-07       sim15.opf   a2-15.legofit
2.062618324e-07       sim16.opf   a2-16.legofit
2.737899195e-07       sim17.opf   a2-17.legofit
2.200061888e-07       sim18.opf   a2-18.legofit
2.787788517e-07       sim19.opf   a2-19.legofit
2.514679774e-07        sim2.opf    a2-2.legofit
1.957039766e-07       sim20.opf   a2-20.legofit
1.960413786e-07       sim21.opf   a2-21.legofit
1.922808068e-07       sim22.opf   a2-22.legofit
5.245407807e-07       sim23.opf   a2-23.legofit
2.622960415e-07       sim24.opf   a2-24.legofit
4.789271767e-07       sim25.opf   a2-25.legofit
3.459524937e-07       sim26.opf   a2-26.legofit
1.802522698e-07       sim27.opf   a2-27.legofit
2.197368688e-07       sim28.opf   a2-28.legofit
2.553795002e-07       sim29.opf   a2-29.legofit
2.355801367e-07        sim3.opf    a2-3.legofit
2.335672304e-07       sim30.opf   a2-30.legofit
2.325515499e-07       sim31.opf   a2-31.legofit
2.096869616e-07       sim32.opf   a2-32.legofit
1.926044104e-07       sim33.opf   a2-33.legofit
3.926801384e-07       sim34.opf   a2-34.legofit
2.843748462e-07       sim35.opf   a2-35.legofit
 2.63153505e-07       sim36.opf   a2-36.legofit
2.519030452e-07       sim37.opf   a2-37.legofit
2.812848171e-07       sim38.opf   a2-38.legofit
2.323723665e-07       sim39.opf   a2-39.legofit
2.409397391e-07        sim4.opf    a2-4.legofit
7.009588266e-07       sim40.opf   a2-40.legofit
2.524176473e-07       sim41.opf   a2-41.legofit
3.152636622e-07       sim42.opf   a2-42.legofit
2.624536864e-07       sim43.opf   a2-43.legofit
 1.95014757e-07       sim44.opf   a2-44.legofit
1.873612868e-07       sim45.opf   a2-45.legofit
2.315575973e-07       sim46.opf   a2-46.legofit
1.937313521e-07       sim47.opf   a2-47.legofit
3.701784227e-07       sim48.opf   a2-48.legofit
1.956634347e-07       sim49.opf   a2-49.legofit
2.502296306e-07        sim5.opf    a2-5.legofit
2.577994756e-07        sim6.opf    a2-6.legofit
2.363159975e-07        sim7.opf    a2-7.legofit
2.057133101e-07        sim8.opf    a2-8.legofit
2.223858837e-07        sim9.opf    a2-9.legofit
```

##### B.9 a2.clic

```
####################################################
```

```
# clic: composite likelihood information criterion #
#                  version 1.81                    #
####################################################

# Program was compiled: Feb  8 2019 15:43:10
# Program was run: Fri Feb  8 15:45:25 2019

#         clic             PtsFile     LegofitFile
   15227155.76         a-sim0-2.pts    a2-0.legofit
   15284394.08         a-sim1-2.pts    a2-1.legofit
   15249975.15        a-sim10-2.pts   a2-10.legofit
   15258098.78        a-sim11-2.pts   a2-11.legofit
   15254485.05        a-sim12-2.pts   a2-12.legofit
   15237601.69        a-sim13-2.pts   a2-13.legofit
   15235499.11        a-sim14-2.pts   a2-14.legofit
   15262919.05        a-sim15-2.pts   a2-15.legofit
   15239043.63        a-sim16-2.pts   a2-16.legofit
   15219332.67        a-sim17-2.pts   a2-17.legofit
   15254238.73        a-sim18-2.pts   a2-18.legofit
   15282190.26        a-sim19-2.pts   a2-19.legofit
   15284694.64         a-sim2-2.pts    a2-2.legofit
    15261621.1        a-sim20-2.pts   a2-20.legofit
   15244942.33        a-sim21-2.pts   a2-21.legofit
   15252230.96        a-sim22-2.pts   a2-22.legofit
   15275233.85        a-sim23-2.pts   a2-23.legofit
    15263782.7        a-sim24-2.pts   a2-24.legofit
    15280545.5        a-sim25-2.pts   a2-25.legofit
   15226111.21        a-sim26-2.pts   a2-26.legofit
   15254065.07        a-sim27-2.pts   a2-27.legofit
   15268215.61        a-sim28-2.pts   a2-28.legofit
   15257601.93        a-sim29-2.pts   a2-29.legofit
    15264118.9         a-sim3-2.pts    a2-3.legofit
   15277340.77        a-sim30-2.pts   a2-30.legofit
   15233652.67        a-sim31-2.pts   a2-31.legofit
   15285148.19        a-sim32-2.pts   a2-32.legofit
   15246164.42        a-sim33-2.pts   a2-33.legofit
   15254437.33        a-sim34-2.pts   a2-34.legofit
    15228382.5        a-sim35-2.pts   a2-35.legofit
   15245678.25        a-sim36-2.pts   a2-36.legofit
   15260903.25        a-sim37-2.pts   a2-37.legofit
   15289393.32        a-sim38-2.pts   a2-38.legofit
   15233628.74        a-sim39-2.pts   a2-39.legofit
    15248980.9         a-sim4-2.pts    a2-4.legofit
   15250088.39        a-sim40-2.pts   a2-40.legofit
   15281272.06        a-sim41-2.pts   a2-41.legofit
   15258074.66        a-sim42-2.pts   a2-42.legofit
    15276895.8        a-sim43-2.pts   a2-43.legofit
   15265381.03        a-sim44-2.pts   a2-44.legofit
```

```
    15197839.46        a-sim45-2.pts   a2-45.legofit
    15252498.16        a-sim46-2.pts   a2-46.legofit
    15242083.73        a-sim47-2.pts   a2-47.legofit
    15235843.93        a-sim48-2.pts   a2-48.legofit
    15228551.07        a-sim49-2.pts   a2-49.legofit
    15239141.11         a-sim5-2.pts    a2-5.legofit
    15232039.77         a-sim6-2.pts    a2-6.legofit
    15248492.16         a-sim7-2.pts    a2-7.legofit
    15278132.82         a-sim8-2.pts    a2-8.legofit
    15265579.38         a-sim9-2.pts    a2-9.legofit
```

### B.10 a2.r

```
library(ggplot2)
library(tidyr)
# Adjust text size. Default is 11
mytheme = theme_get()
mytheme$text$size = 20
theme_set(mytheme)

wide <- read.table("a2.flat", header=T)

# parameter names
parnames <- c("mN", "mD", "Txy", "Tnd", "Ta", "Td",
              "twoNxynd", "twoNxy", "twoNnd", "twoNn", "twoNd")

wide <- wide[,parnames]

# parameter labels
lbl <- expression("m"["N"], "m"["D"], "T"["XY"], "T"["ND"], "T"["A"], "T"["D"],
    "2N"["XYND"], "2N"["XY"], "2N"["ND"], "2N"["N"], "2N"["D"])

kappa(cor(wide))
pairs(wide, labels=lbl)

pdf("msp-a2-pairs.pdf")
pairs(wide, labels=lbl)
dev.off()
```

##### B.11 b.lgo

```
# (grep ^# a.lgo; pclgo a.lgo a2-*.legofit; grep -v ^# a.lgo | egrep -v
# "\<free\>") > b.lgo
# Model: ((X,Y), (N,D)), migration: N -> Y, D -> Y
# This file is like true.lgo but the free parameter values have been
# arbitrarily moved to incorrect values so that legofit will have some
# work to do.
# PCA calculated with gsl_linalg_SV_decomp
```

```
# Fraction of variance:
#      pc1      pc2      pc3      pc4      pc5      pc6      pc7      pc8
#  0.55901  0.22252  0.13678  0.03296  0.02182  0.01016  0.00904  0.00619
#      pc9     pc10     pc11
#  0.00133  0.00011  0.00007
param free [    -4,      4] pc1 = 0
param free [    -2,      2] pc2 = 0
param free [    -2,      2] pc3 = 0
param free [    -2,      2] pc4 = 0
param free [    -2,      2] pc5 = 0
param free [    -2,      2] pc6 = 0
param free [    -1,      1] pc7 = 0
param free [    -1,      1] pc8 = 0
param free [    -1,      1] pc9 = 0
param free [  -0.5,    0.5] pc10 = 0
param free [  -0.5,    0.5] pc11 = 0
time constrained Tnd = 15911.022 + 247.38646*pc1 + 121.891*pc2 +
    123.7284*pc3 - 34.949662*pc4 - 48.53957*pc5 + 78.332907*pc6 -
    66.3194*pc7 - 209.04369*pc8 + 57.980567*pc9 + 526.8714*pc10 -
    105.40331*pc11
time constrained Txy = 6861.6542 + 823.19668*pc1 + 567.87533*pc2 +
    162.20725*pc3 + 216.13627*pc4 + 143.07624*pc5 - 369.48603*pc6 +
    135.08397*pc7 + 953.72375*pc8 + 186.6496*pc9 - 428.78426*pc10 -
    1650.1013*pc11
time constrained Td = 1805.285 + 3.5283167*pc1 - 42.411078*pc2 +
    66.651041*pc3 + 42.781523*pc4 - 66.999364*pc5 - 11.779426*pc6 +
    12.232257*pc7 + 5.4324914*pc8 - 4.9049812*pc9 - 4.6563794*pc10 +
    0.94483801*pc11
time constrained Ta = 1821.7234 + 1.3646556*pc1 - 21.359967*pc2 +
    35.38505*pc3 - 21.510298*pc4 + 35.18434*pc5 + 3.1387016*pc6 -
    0.74743938*pc7 + 6.5075683*pc8 - 4.1822283*pc9 + 0.19043585*pc10 -
    0.48381102*pc11
twoN constrained twoNnd = 4556.4996 - 115.48227*pc1 - 53.86141*pc2 -
    61.76295*pc3 + 18.853564*pc4 + 6.5344732*pc5 - 66.585286*pc6 +
    27.709276*pc7 + 131.15742*pc8 - 165.52022*pc9 + 157.50935*pc10 -
    45.850625*pc11
twoN constrained twoNn = 7368.7528 + 534.39557*pc1 - 618.18898*pc2 -
    433.04216*pc3 + 724.90564*pc4 + 661.13922*pc5 - 467.21813*pc6 +
    997.40853*pc7 - 620.48085*pc8 + 11.449352*pc9 + 46.439577*pc10 -
    19.569022*pc11
twoN constrained twoNd = 8033.262 - 739.6403*pc1 + 793.58728*pc2 +
    624.83955*pc3 + 1370.9644*pc4 + 864.0819*pc5 - 877.93833*pc6 -
    1153.3321*pc7 - 538.2864*pc8 - 82.445814*pc9 - 29.739809*pc10 +
    9.4925392*pc11
twoN constrained twoNxy = 37875.408 - 1812.3968*pc1 - 1244.4373*pc2 -
    333.73719*pc3 - 535.1146*pc4 - 418.52206*pc5 + 1136.101*pc6 -
    414.62503*pc7 - 2644.9723*pc8 - 679.7313*pc9 - 689.95397*pc10 -
    3186.291*pc11
```

```
twoN constrained twoNxynd = 67325.848 + 693.706*pc1 + 514.18721*pc2 +
    176.84976*pc3 - 65.126123*pc4 - 64.775154*pc5 + 199.23202*pc6 +
    23.77747*pc7 - 364.79298*pc8 - 1580.8074*pc9 - 451.62602*pc10 +
    211.30396*pc11
mixFrac constrained mN = 0.044541498 + 0.0012489876*pc1 -
    0.0017511182*pc2 - 0.001037823*pc3 + 0.0018636076*pc4 +
    0.00059977129*pc5 + 0.0024179158*pc6 - 0.0021985549*pc7 +
    0.0010036654*pc8 - 0.00020303422*pc9 + 3.3897379e-05*pc10 +
    4.1465522e-05*pc11
mixFrac constrained mD = 0.03076842 - 0.0013156222*pc1 +
    0.0015223186*pc2 + 0.0008878441*pc3 + 0.0013664654*pc4 +
    0.00064836698*pc5 + 0.0027362886*pc6 + 0.0021675192*pc7 +
    0.00046269859*pc8 + 0.00013389589*pc9 + 0.0002374218*pc10 -
    2.433369e-05*pc11
time fixed   zero  = 0
twoN fixed   one   = 1
time fixed   Txynd = 25920   # \citet[table~S12.2, p.~90]{Li:N-505-43-S88}
time fixed     TmN = 1897    # \citep[table~2]{Sankararaman:PLO-8-e1002947}
time fixed     TmD = 1       # arbitrary
segment x    t=zero   twoN=one     samples=1
segment y    t=zero   twoN=one     samples=1
segment n    t=Ta     twoN=twoNn   samples=1
segment n2   t=TmN    twoN=twoNn
segment d0   t=TmD    twoN=one
segment d    t=Td     twoN=twoNd   samples=1
segment y1   t=TmD    twoN=one
segment y2   t=TmN    twoN=one
segment nd   t=Tnd    twoN=twoNnd
segment xy   t=Txy    twoN=twoNxy
segment xynd t=Txynd  twoN=twoNxynd
mix    y  from y1 + mD * d0
mix    y1 from y2 + mN * n2
derive x  from xy
derive y2 from xy
derive n  from n2
derive n2 from nd
derive d0 from d
derive d  from nd
derive xy from xynd
derive nd from xynd
```

##### B.12 c.lgo

```
# (grep ^# a.lgo; pclgo --tol 0.001 a.lgo a2-*.legofit; grep -v ^# a.lgo |
# egrep -v "\<free\>") > c.lgo
# Model: ((X,Y), (N,D)), migration: N -> Y, D -> Y
# This file is like true.lgo but the free parameter values have been
# arbitrarily moved to incorrect values so that legofit will have some
```

```
# work to do.
# PCA calculated with gsl_linalg_SV_decomp
# Fraction of variance:
#      pc1      pc2      pc3      pc4      pc5      pc6      pc7      pc8
#  0.55901  0.22252  0.13678  0.03296  0.02182  0.01016  0.00904  0.00619
#      pc9     pc10     pc11
#  0.00133  0.00011  0.00007
param free [    -4,      4] pc1 = 0
param free [    -2,      2] pc2 = 0
param free [    -2,      2] pc3 = 0
param free [    -2,      2] pc4 = 0
param free [    -2,      2] pc5 = 0
param free [    -2,      2] pc6 = 0
param free [    -1,      1] pc7 = 0
param free [    -1,      1] pc8 = 0
param free [    -1,      1] pc9 = 0
time constrained Tnd = 15911.022 + 247.38646*pc1 + 121.891*pc2 +
    123.7284*pc3 - 34.949662*pc4 - 48.53957*pc5 + 78.332907*pc6 -
    66.3194*pc7 - 209.04369*pc8 + 57.980567*pc9
time constrained Txy = 6861.6542 + 823.19668*pc1 + 567.87533*pc2 +
    162.20725*pc3 + 216.13627*pc4 + 143.07624*pc5 - 369.48603*pc6 +
    135.08397*pc7 + 953.72375*pc8 + 186.6496*pc9
time constrained Td = 1805.285 + 3.5283167*pc1 - 42.411078*pc2 +
    66.651041*pc3 + 42.781523*pc4 - 66.999364*pc5 - 11.779426*pc6 +
    12.232257*pc7 + 5.4324914*pc8 - 4.9049812*pc9
time constrained Ta = 1821.7234 + 1.3646556*pc1 - 21.359967*pc2 +
    35.38505*pc3 - 21.510298*pc4 + 35.18434*pc5 + 3.1387016*pc6 -
    0.74743938*pc7 + 6.5075683*pc8 - 4.1822283*pc9
twoN constrained twoNnd = 4556.4996 - 115.48227*pc1 - 53.86141*pc2 -
    61.76295*pc3 + 18.853564*pc4 + 6.5344732*pc5 - 66.585286*pc6 +
    27.709276*pc7 + 131.15742*pc8 - 165.52022*pc9
twoN constrained twoNn = 7368.7528 + 534.39557*pc1 - 618.18898*pc2 -
    433.04216*pc3 + 724.90564*pc4 + 661.13922*pc5 - 467.21813*pc6 +
    997.40853*pc7 - 620.48085*pc8 + 11.449352*pc9
twoN constrained twoNd = 8033.262 - 739.6403*pc1 + 793.58728*pc2 +
    624.83955*pc3 + 1370.9644*pc4 + 864.0819*pc5 - 877.93833*pc6 -
    1153.3321*pc7 - 538.2864*pc8 - 82.445814*pc9
twoN constrained twoNxy = 37875.408 - 1812.3968*pc1 - 1244.4373*pc2 -
    333.73719*pc3 - 535.1146*pc4 - 418.52206*pc5 + 1136.101*pc6 -
    414.62503*pc7 - 2644.9723*pc8 - 679.7313*pc9
twoN constrained twoNxynd = 67325.848 + 693.706*pc1 + 514.18721*pc2 +
    176.84976*pc3 - 65.126123*pc4 - 64.775154*pc5 + 199.23202*pc6 +
    23.77747*pc7 - 364.79298*pc8 - 1580.8074*pc9
mixFrac constrained mN = 0.044541498 + 0.0012489876*pc1 -
    0.0017511182*pc2 - 0.001037823*pc3 + 0.0018636076*pc4 +
    0.00059977129*pc5 + 0.0024179158*pc6 - 0.0021985549*pc7 +
    0.0010036654*pc8 - 0.00020303422*pc9
mixFrac constrained mD = 0.03076842 - 0.0013156222*pc1 +
```

```
    0.0015223186*pc2 + 0.0008878441*pc3 + 0.0013664654*pc4 +
    0.00064836698*pc5 + 0.0027362886*pc6 + 0.0021675192*pc7 +
    0.00046269859*pc8 + 0.00013389589*pc9
time fixed   zero  = 0
twoN fixed   one   = 1
time fixed   Txynd = 25920   # \citet[table~S12.2, p.~90]{Li:N-505-43-S88}
time fixed     TmN = 1897    # \citep[table~2]{Sankararaman:PLO-8-e1002947}
time fixed     TmD = 1       # arbitrary
segment x    t=zero   twoN=one     samples=1
segment y    t=zero   twoN=one     samples=1
segment n    t=Ta     twoN=twoNn   samples=1
segment n2   t=TmN    twoN=twoNn
segment d0   t=TmD    twoN=one
segment d    t=Td     twoN=twoNd   samples=1
segment y1   t=TmD    twoN=one
segment y2   t=TmN    twoN=one
segment nd   t=Tnd    twoN=twoNnd
segment xy   t=Txy    twoN=twoNxy
segment xynd t=Txynd  twoN=twoNxynd
mix    y  from y1 + mD * d0
mix    y1 from y2 + mN * n2
derive x  from xy
derive y2 from xy
derive n  from n2
derive n2 from nd
derive d0 from d
derive d  from nd
derive xy from xynd
derive nd from xynd
```

#### B.13 bma.r

```
library(ggplot2)
library(tidyr)
# Adjust text size. Default is 11
mytheme = theme_get()
mytheme$text$size = 25
theme_set(mytheme)

wide <- read.table("a-b-c.bma", header=T)

# parameter names
parnames <- c("mN", "mD", "Txy", "Tnd", "Ta", "Td",
              "twoNxynd", "twoNxy", "twoNnd", "twoNn", "twoNd")

wide <- wide[,parnames]

# parameter labels
```

```
lbl <- expression("m"["N"], "m"["D"], "T"["XY"], "T"["ND"], "T"["A"], "T"["D"],
    "2N"["XYND"], "2N"["XY"], "2N"["ND"], "2N"["N"], "2N"["D"])

# Migration rates
mpar <- c("mN", "mD")
mdat <- subset(wide, select=mpar)
mdat$msum <- mdat$mN + mdat$mD
mdat$mdif <- mdat$mN - mdat$mD
mlbl <- expression("m"["N"], "m"["D"], "m"["N"]+"m"["D"], "m"["N"]-"m"["D"])
mdat <- gather(mdat, par,value, mN, mD, msum, mdif)
mdat$tru <- rep(NA, nrow(mdat))
mdat$tru[mdat$par == "mN"] <- 0.05
mdat$tru[mdat$par == "mD"] <- 0.025
mdat$tru[mdat$par == "msum"] <- 0.075
mdat$tru[mdat$par == "mdif"] <- 0.025
mpar <- c("mN", "mD", "msum", "mdif")
mdat$par <- ordered(mdat$par, levels=rev(mpar))

# Plot migration rate
ggplot(mdat, aes(x=par, y=value)) +
    scale_x_discrete(labels=rev(mlbl)) +
    geom_violin(scale="width") +
    geom_point(aes(par, tru), shape=4, size=4, color="red") +
    ylab("Admixture Fraction") +
    scale_y_continuous(limits=c(0,0.08)) +
    theme(aspect.ratio=0.2,
          axis.title.y = element_blank()) +
    coord_flip()
ggsave("msp-bma-mviol.pdf")

ggplot(mdat, aes(value, par)) +
    xlab("Admixture Fraction") +
    theme(aspect.ratio=0.2,
          axis.title.y = element_blank()) +
    scale_x_continuous(limits=c(0,0.091)) +
    scale_y_discrete(labels=rev(mlbl)) +
    geom_jitter(height=0.2, width=0, shape=1, alpha=0.5, size=4, color="blue") +
    geom_point(mapping=aes(tru, par), shape=4, size=4, color="red")
ggsave("msp-bma-mdot.pdf")

# Time parameters
tpar <- c("Txy", "Tnd", "Ta", "Td")
tlbl <- expression("T"["XY"], "T"["ND"], "T"["A"], "T"["D"])
tdat <- subset(wide, select=tpar)
tdat <- gather(tdat, par,value)
tdat$tru <- rep(NA, nrow(tdat))
tdat$tru[tdat$par == "Txy"] <- 3788
tdat$tru[tdat$par == "Tnd"] <- 15000
```

```
tdat$tru[tdat$par == "Ta"] <- 1760
tdat$tru[tdat$par == "Td"] <- 1734
tdat$par <- ordered(tdat$par, levels=tpar)

# Plot times
ggplot(tdat, aes(x=par, y=value)) +
    scale_x_discrete(labels=tlbl) +
    geom_violin(scale="width") +
    geom_point(aes(par, tru), shape=4, size=4, color="red") +
    ylab("Generations") +
    theme(aspect.ratio=0.3,
          panel.grid.minor = element_blank(),
          axis.title.y = element_blank()) +
    coord_flip()
ggsave("msp-bma-tviol.pdf")

ggplot(tdat, aes(29*value/1000, par)) +
    xlab("Thousands of Years") +
    theme(aspect.ratio=0.2,
          axis.title.y = element_blank()) +
    scale_y_discrete(labels=tlbl) +
    geom_jitter(height=0.2, width=0, shape=1, alpha=0.5, size=4, color="blue") +
    geom_point(mapping=aes(29*tru/1000, par), shape=4, size=4, color="red")
ggsave("msp-bma-tdot.pdf")

# Pop sizes
npar <- c("twoNxynd", "twoNxy", "twoNnd", "twoNn", "twoNd")
nlbl <- expression("2N"["XYND"], "2N"["XY"], "2N"["ND"], "2N"["N"],
    "2N"["D"])
ndat <- subset(wide, select=npar)
ndat <- gather(ndat, par,value)
ndat$tru <- rep(NA, nrow(ndat))
ndat$tru[ndat$par == "twoNxynd"] <- 64964.1
ndat$tru[ndat$par == "twoNxy"] <- 44869.2
ndat$tru[ndat$par == "twoNnd"] <- 5000.0
ndat$tru[ndat$par == "twoNn"] <- 9756.8
ndat$tru[ndat$par == "twoNd"] <- 5000
ndat$par <- ordered(ndat$par, levels=npar)

# Plot pop sizes
ggplot(ndat, aes(x=par, y=value)) +
    scale_x_discrete(labels=nlbl) +
    geom_violin(scale="width") +
    geom_point(aes(par, tru), shape=4, size=4, color="red") +
#    ylim(15000, 25000) +
    ylab("Haploid Population Size (2N)") +
    theme(aspect.ratio=0.3,
          axis.title.y = element_blank()) +
```

```
    coord_flip()
ggsave("msp-bma-nviol.pdf")

ggplot(ndat, aes(value, par)) +
    xlab("Haploid Population Size (2N)") +
    theme(aspect.ratio=0.3,
          axis.title.y = element_blank()) +
    scale_y_discrete(labels=nlbl) +
    geom_jitter(height=0.2, width=0, shape=1, alpha=0.5, size=4, color="blue") +
    geom_point(mapping=aes(tru, par), shape=4, size=4, color="red")
ggsave("msp-bma-ndot.pdf")
```

##### B.14 nma2resid.r

##### B.15 neaden.py

```
#!/usr/bin/python

###
#@file neaden.py
#@page neaden
#@brief Calculate "nea" and "den" (aka R_Neandertal)
#
# Names of input files should be listed on the command line. Within
# each input file, blank lines and lines beginning with '#' are
# ignored. Other lines are split into whitespace-separated fields. The
# first field is interpreted as the label of a site pattern, the
# second should be proportional to the frequency of that site
# pattern. The program normalizes frequencies so that they sum to 1.
# neaden writes a header line: "nea den". After that, each line
# contains two values, nea and den, which estimate Neanderthal and
# Denisovan admixture. These estimators were defined in supplementary
# note 11 of Meyer et al (2012, Science, 328(5979):710-722).

from math import log
import sys

def usage():
    print "usage: neaden.py inputfile1 [inputfile2 ...]"
    print "    In input site pattern labels, x=African, y=Eurasian,"
    print "    n=Neanderthal, and d=Denisovan"
    exit(1)

def process_file(fname):
    f = open(fname)

    pr = {}
    s = 0.0
```

```
    # Build map of keys to probabilities
    for line in f:
        line = line.strip()
        if len(line)==0 or line[0] == '#':
            continue
        line = line.split()
        key = line[0]
        x = float(line[1])
        pr[key] = x
        s += x

    f.close()

    # Normalize probabilities
    for key in pr.keys():
        pr[key] /= s

    # Eqn 5 in Supp Note 11, Meyer et al 2012
    nea = pr["y:d"] + pr["y:n:d"] - pr["x:d"] - pr["x:n:d"]
    nea /= pr["n:d"] + pr["y:n:d"] - pr["x:d"] - pr["x:y:d"]

    # Eqn 8 in Supp Note 11, Meyer et al 2012
    den = pr["y:d"] + pr["y:n:d"] - pr["x:n"] - pr["x:n:d"]
    den /= pr["n:d"] + pr["y:n:d"] - pr["x:n"] - pr["x:y:n"]

    return (nea, den)

if len(sys.argv) < 2:
    usage()

# abort with usage message if there are any flag arguments
for arg in sys.argv[1:]:
    if arg[0] == '-':
        usage()

# Print a line of output for each input file. Each line consists of
# two values: an estimate of Neanderthal admixture and one of
# Denisovan admixture.
print "%10s %10s" % ("nea", "den")
for fname in sys.argv[1:]:
    nea, den = process_file(fname)
    print "%10.8f %10.8f" % (nea, den)
```

##### B.16 neaden.r

```
library(ggplot2)
library(tidyr)
# Adjust text size. Default is 11
mytheme = theme_get()
mytheme$text$size = 25
theme_set(mytheme)

wide <- read.table("neaden.out", header=T)
wide$TreeMix <- scan("../treemix/treemix-nea.txt")

mdat <- gather(wide, par,value)
mdat$tru <- rep(NA, nrow(mdat))
mdat$tru[mdat$par == "nea"] <- 0.05
mdat$tru[mdat$par == "den"] <- 0.025
mdat$tru[mdat$par == "TreeMix"] <- 0.05
mdat$par <- ordered(mdat$par, levels=rev(c("nea", "den", "TreeMix")))

# Expected values calculated as ./neaden.py ../legosim/true.patfrq:
Enea <- 0.08137593
Eden <- 0.08183959

mdat$expct <- rep(NA, nrow(mdat))
mdat$expct[mdat$par == "nea"] <- Enea
mdat$expct[mdat$par == "den"] <- Eden

# Plot migration rate
ggplot(mdat, aes(value, par)) +
    xlab("Admixture Fraction") +
    scale_x_continuous(limits=c(0,0.091)) +
    theme(aspect.ratio=0.2,
          axis.title.y = element_blank()) +
    geom_jitter(height=0.2, width=0, shape=1, alpha=0.5, size=4, color="blue") +
    geom_point(mapping=aes(tru, par), shape=4, size=4, color="red") +
    geom_point(mapping=aes(expct, par), shape=2, size=4, color="black")
ggsave("neaden.pdf")
```

## C Ms

### C.1 ms.py

```
#!/usr/bin/python
# Model ((X,Y),(N,D)) with gene flow from N -> Y
import os, sys, time
from math import expm1
```

```
def usage():
    print "Usage: ms.py [options]"
    print "  where options may include:"
    print "  -r or --run: run ms. By default, command is printed"
    print "               but not executed."
    sys.exit(1)

runprogram = False
for arg in sys.argv[1:]:
    if arg == "-r" or arg == "--run":
        runprogram = True
    else:
usage()

# Populations are X, Y, N, D

# time parameters in generations
Txynd = 25920.0
Tnd = 15000.0
Txy = 3788.0
Td = 1734.0    # age of Denisova fossil
Ta = 1760.0    # age of Altai fossil
TmN = 1897.0   # time of Neanderthal admixture
TmD = Td-1.0   # time of Denisovan admixture

# haploid population sizes
twoN0 = 64964.1  # ancestral population
twoNxy = 44869.2
twoNnd = 5000.0
twoNn = 9756.8
twoNd = 5000.0
twoNx = 20000.0 # modern Africa
twoNy = 20000.0 # modern Europe

# admixture
mN = 0.05
mD = 0.025

nreps = 1000         # number of chromosomes
twoNsmp = 2          # number of modern haploid samples
basepairs = int(2e6) # number of nucleotides per chromosome
u_per_site = 1.4e-8  # mutation
u_per_seq = u_per_site * basepairs

# Recombination rate.  recomb is the probability of a cross-over
# between sites at opposite ends of the simulated sequence.
c = 1e-8 # rate per base pair per generation
recomb = c * (basepairs-1)
```

```
# Random number seeds. s1 is a mixture of the current time and the
# process id. ms wants three seeds, so I generate the other two by
# fiddling with the bits of s1. Specifically, I shift the even bits
# left and the odd ones right to make s2, then do the opposite shift
# to make s3. There is no theory behind this. I'm just making up
# numbers.
s1 = int(time.time()) ^ os.getpid()
even_bits = s1 & 0xAAAAAAAA
odd_bits = s1 & 0x55555555
s2 = (even_bits << 1) | (odd_bits >> 1)
s3 = (odd_bits << 1) | (even_bits >> 1)

# construct command string
cmd = "ms %d %d \\\n" % (twoNsmp, nreps)
cmd += " -t %g \\\n" % (u_per_seq * 2 * twoN0) # theta

# random number seeds
cmd += " -seeds %d %d %d \\\n" % (s1, s2, s3)

# Argument of -r is 2*twoN0*C basepairs, where C =
# recombination*(basepairs-1) is recombination rate between sites at
# opposite ends of the sequence and basepairs is the number of base
# pairs in the sequence.
cmd += " -r %g %d \\\n" % (recomb * 2 * twoN0, basepairs)

# npops and haploid samples/pop at time 0
# Populations are 1=X (Africa), 2=Y (Eurasia), 3=N (Neanderthal), and
# 4=D (Denisovan).
cmd += " -I 4 1 1 0 0 \\\n"

# population sizes
cmd += " -n 1 %g \\\n" % (twoNx/twoN0)  # Africa
cmd += " -n 2 %g \\\n" % (twoNy/twoN0)  # Europe
cmd += " -n 3 %g \\\n" % (twoNn/twoN0)  # Neanderthal
cmd += " -n 4 %g \\\n" % (twoNd/twoN0)  # Denisova
cmd += " -en %g 1 %g \\\n" % (Txy/(2*twoN0), twoNxy/twoN0) # XY
cmd += " -en %g 1 %g \\\n" % (Txynd/(2*twoN0), 1.0)        # XYND
cmd += " -en %g 3 %g \\\n" % (Tnd/(2*twoN0), twoNnd/twoN0) # ND

# Fossils
cmd += " -eA %g 3 1 \\\n" % (Ta/(2*twoN0)) # Altai
cmd += " -eA %g 4 1 \\\n" % (Td/(2*twoN0)) # Denisova

# Each gene flow event takes two steps. First, the -es argument moves
# a fraction of the lineages in 2 into a newly-created
# population. Then, an -ej argument moves these into the population
# we're aiming at.
```

```
# D->Y gene flow. First step generates population 5
cmd += " -es %g 2 %g \\\n" % (TmD/(2*twoN0), 1-mD)
# Second step merges 5 into 4.
cmd += " -ej %g 5 4 \\\n" % (TmD/(2*twoN0))

## N->Y gene flow. First step generates population 6.
cmd += " -es %g 2 %g \\\n" % (TmN/(2*twoN0), 1-mN)
# Second step merges 6 into 3.
cmd += " -ej %g 6 3 \\\n" % (TmN/(2*twoN0))

# population splits
cmd += " -ej %g 2 1 \\\n" % (Txy/(2*twoN0)) # X-Y split
cmd += " -ej %g 4 3 \\\n" % (Tnd/(2*twoN0)) # N-D split
cmd += " -ej %g 3 1\n" % (Txynd/(2*twoN0)) # XY-ND split

if runprogram:
    os.system(cmd)
else:
    print "Dry run:", cmd
```

##### C.2 sim.sh

```
# ms.py is a Python script that runs ms.  ms2sim and simpat are part
# of the Legofit package. ms2sim converts ms output into sim
# format. Simpat reads this reformatted output and tabulates site
# patterns.
ofile=sim${1}.opf
efile=sim${1}.err
python ms.py -r | ms2sim x:0 y:1 d:3 n:2 | simpat 1>${ofile} 2>${efile}
```

#### D Scrm

##### D.1 scrm.py

```
#!/usr/bin/python
# Model ((X,Y),(N,D)) with gene flow from N -> Y
import os, sys
from math import expm1

def usage():
    print "Usage: scrm.py [options]"
    print "  where options may include:"
    print "  -r or --run: run scrm. By default, command is printed"
    print "               but not executed."
    sys.exit(1)
```

```
runprogram = False
for arg in sys.argv[1:]:
    if arg == "-r" or arg == "--run":
        runprogram = True
    else:
usage()

# Parameters
# Parameter values from mig.lgo
# Populations are X, Y, N, D

# time parameters in generations
Txynd = 25920
Tnd = 15000
Txy = 3788
Td = 1734      # age of Denisova fossil
Ta = 1760      # age of Altai fossil
TmN = 1897     # time of Neanderthal admixture
TmD = Td-1     # time of Denisovan admixture

# haploid population sizes
twoN0 = 64964.1  # ancestral population
twoNxy = 44869.2
twoNnd = 5000
twoNn = 9756.8
twoNd = 5000
twoNx = 20000.0 # modern Africa
twoNy = 20000.0 # modern Europe

# admixture
mN = 0.05
mD = 0.025

nreps = 1000      # number of chromosomes
twoNsmp = 4       # total number of haploid samples
basepairs = 2e6   # number of nucleotides per chromosome
u_per_site = 1.4e-8 # mutation
u_per_seq = u_per_site * basepairs

# Recombination rate.  recomb is the probability of recombination
# between sites at opposite ends of the simulated sequence.
c = 1e-8 # rate per base pair per generation
recomb = -expm1(-2.0*c*(basepairs-1))/2.0 # Haldane's mapping rule

# construct command string
cmd = "scrm %s %s \\\n" % (twoNsmp, nreps)
cmd += " -l 500r \\\n" # controls accuracy of approximation
cmd += " -t %g \\\n" % (u_per_seq * 2 * twoN0) # theta
```

```
cmd += " -r %g %g \\\n" % (recomb * 2 * twoN0, basepairs)
cmd += " -transpose-segsites \\\n"   # mutations in rows, h'types in cols
cmd += " -SC abs \\\n"               # sequence positions in base pairs

# npops and haploid samples/pop at time 0
cmd += " -I 4 1 1 0 0 \\\n"

# Fossils
cmd += " -eI %g 0 0 1 0 \\\n" % (Ta/(2*twoN0)) # Altai
cmd += " -eI %g 0 0 0 1 \\\n" % (Td/(2*twoN0)) # Denisova

# population sizes
cmd += " -n 1 %g \\\n" % (twoNx/twoN0)  # Africa
cmd += " -n 2 %g \\\n" % (twoNy/twoN0)  # Europe
cmd += " -n 3 %g \\\n" % (twoNn/twoN0)  # Neanderthal
cmd += " -n 4 %g \\\n" % (twoNd/twoN0)  # Denisova
cmd += " -en %g 1 %g \\\n" % (Txy/(2*twoN0), twoNxy/twoN0) # XY
cmd += " -en %g 1 %g \\\n" % (Txynd/(2*twoN0), 1.0)        # XYND
cmd += " -en %g 3 %g \\\n" % (Tnd/(2*twoN0), twoNnd/twoN0) # ND

# population splits
cmd += " -ej %g 3 1 \\\n" % (Txynd/(2*twoN0)) # XY-ND split
cmd += " -ej %g 2 1 \\\n" % (Txy/(2*twoN0)) # X-Y split
cmd += " -ej %g 4 3 \\\n" % (Tnd/(2*twoN0)) # N-D split

# gene flow
cmd += " -eps %g 2 4 %g \\\n" % (TmD/(2*twoN0), 1-mD) # D->Y gene flow
cmd += " -eps %g 2 3 %g \\\n" % (TmN/(2*twoN0), 1-mN) # N->Y gene flow

if runprogram:
    os.system(cmd)
else:
    print "Dry run:", cmd
```

#### D.2 sim.slr

```
#!/bin/bash
#SBATCH -J sim
#SBATCH --account=rogersa-kp
#SBATCH --partition=rogersa-kp
#SBATCH --time=24:00:00
#SBATCH --nodes 1
#SBATCH --ntasks 1
#SBATCH -o sim%a.opf # stdout
#SBATCH -e sim%a.err # stderr
```

```
# Execute this with "sbatch --array=0-49 sim.slr".  This will
# launch a job for each simulation replicate.

# scrm.py is a Python script that runs scrm. scrmpat is part of the
# Legofit package. It reads scrm output and tabulates site patterns.

time python scrm.py -r | scrmpat x y n d
```

### E TreeMix

#### E.1 README.md

```
# Experiment using TreeMix

treemix.py does the same simulation as ../msp/msp.py, but writes the
output in treemix format. treemix.sh takes one argument, which
specifies the index of the current simulation. It writes files with
names like sim0.xxx, where 0 is the index of the current simulation.
To generate 50 sets of simulated data:

    seq 0 49 | xargs -n 1 -P 32 bash treemix.sh

The migration rates are in files with names like sim*.treeout.gz. To
get the N->Y migration weights:

    zcat sim*.treeout.gz | grep -v \( | grep "y:" | grep "n:" | awk \
    '{print $1}' > treemix-nea.txt

All 50 output files report a N->Y migration event. None of them report
a D->Y migration event, and only one of them found Y->D.

I found no zeros in the Newick trees in sim*.treeout.gz, so I didn't
use the -noss option.

The data in treemix-nea.txt are graphed in ../msp/neaden.r.
```

#### E.2 treemix.py

```
#!/usr/bin/python3
import msprime
import os, sys, time

def usage():
    print("Usage: ./treemix.py [options]")
    print("  where options may include:")
    print("  -r or --run: run simulation. Default: run")
    print("               DemographyDebugger")
    sys.exit(1)
```

```
do_simulation = False
for arg in sys.argv[1:]:
    if arg == "-r" or arg == "--run":
        do_simulation = True
    else:
        usage()

# time parameters in generations
Txynd = 25920
Tnd = 15000
Txy = 3788
Td = 1734      # age of Denisova fossil
Ta = 1760      # age of Altai fossil
TmN = 1897     # time of Neanderthal admixture
TmD = Td-1     # time of Denisovan admixture

# haploid population sizes
Nxynd = 64964.1/2.0  # ancestral population
Nxy = 44869.2/2.0
Nnd = 5000/2.0
Nn = 9756.8/2.0
Nd = 5000/2.0
Nx = 20000/2.0 # modern Africa
Ny = 20000/2.0 # modern Europe

# admixture
mN = 0.05
mD = 0.025

nchromosomes = 1000      # number of chromosomes
basepairs = 2e6   # number of nucleotides per chromosome
u_per_site = 1.4e-8 # mutation

# Recombination rate.  recomb is the probability of recombination
# between sites at opposite ends of the simulated sequence.
c = 1e-8 # rate per base pair per generation

# One haploid sample from each of 4 populations: two modern (X,Y),
# and two archaic (N,D).
samples = [
    msprime.Sample(population=0, time=0),  # population X
    msprime.Sample(population=1, time=0),  # population Y
    msprime.Sample(population=2, time=Ta), # population N
    msprime.Sample(population=3, time=Td)  # population D
]

lbl = ("x", "y", "n", "d")
```

```
npops = len(lbl)

# associate sample sizes with population labels
sampsize = {}
for s in lbl:
    sampsize[s] = 0

for samp in samples:
    ndx = samp.population
    assert ndx < len(lbl)
    assert lbl[ndx] in sampsize
    sampsize[lbl[ndx]] = 1

for s in lbl:
    if not (sampsize[s] > 0):
        print("Population %s has no samples" % s, file=sys.stderr)
        sys.exit(1)

# Population configurations. No sample sizes are listed here, because
# those are specified above in "samples".
popconf = [
    msprime.PopulationConfiguration(initial_size=Nx),
    msprime.PopulationConfiguration(initial_size=Ny),
    msprime.PopulationConfiguration(initial_size=Nn),
    msprime.PopulationConfiguration(initial_size=Nd)
]

events = [
    msprime.MassMigration(
        time=TmD,
        source=1,
        dest=3,
        proportion=mD), # D->Y gene flow
    msprime.MassMigration(
        time=TmN,
        source=1,
        dest=2,
        proportion=mN), # N->Y gene flow
    msprime.MassMigration(
        time=Txy,
        source=1,
        dest=0,
        proportion=1.0), # X-Y split
    msprime.PopulationParametersChange(
        time=Txy,
        initial_size=Nxy,
        population_id=0),
    msprime.MassMigration(
```

```
        time=Tnd,
        source=3,
        dest=2,
        proportion=1.0), # N-D split
    msprime.PopulationParametersChange(
        time=Tnd,
        initial_size=Nnd,
        population_id=2),
    msprime.MassMigration(
        time=Txynd,
        source=2,
        dest=0,
        proportion=1.0), # XY-ND split
    msprime.PopulationParametersChange(
        time=Txynd,
        initial_size=Nxynd,
        population_id=0)
]

if do_simulation:
    # run simulation
    seed = int(time.time()) ^ os.getpid()
    sim = msprime.simulate(samples = samples,
                           population_configurations = popconf,
                           demographic_events = events,
                           length = basepairs,
                           recombination_rate = c,
                           mutation_rate = u_per_site,
                           num_replicates = nchromosomes,
                           random_seed = seed)

    # header
    for s in lbl:
        print(s, end=" ")
    print("o") # outgroup

    # This code assumes that each population has
    # haploid sample size 1. It will break if sample
    # sizes are larger.
    for i, chromosome in enumerate(sim):
        for variant in chromosome.variants():
            for g in variant.genotypes:
                print("%d,%d" % (g, 1-g), end=" ")
            print("0,1") # outgroup carries ancestral allele at each site

else:
    # Run demography debugger and quit
    dd = msprime.DemographyDebugger(
```

```
        population_configurations=popconf,
        demographic_events=events)
    dd.print_history()
    print("Use \"./sim -r\" to run simulation")
```

##### E.3 treemix.sh

```
#!/bin/bash

if [[ $# -ne 1 ]]
then
echo "usage: bash treemix.sh index_of_current_simulation"
exit 1
fi

# basename of output files depends on argument $1
oname=sim$1
tmpfile=tmp$1.gz

./treemix.py -r | gzip -c > $tmpfile

treemix -i $tmpfile -k 500 -root o -m 2 -o $oname

rm $tmpfile
```

[3] Matthias Meyer, Martin Kircher, Marie-Theres Gansauge, Heng Li, Fernando Racimo, Swapan Mallick, Joshua G Schraiber, Flora Jay, Kay Prüfer, Cesare de Filippo, Peter H. Sudmant, Can Alkan, Qiaomei Fu, Ron Do, Nadin Rohland, Arti Tandon, Michael Siebauer, Richard E. Green, Katarzyna Bryc, Adrian W. Briggs, Udo Stenzel, Jesse Dabney, Jay Shendure, Jacob Kitzman, Michael F. Hammer, Michael V. Shunkov, Anatoli P. Derevianko, Nick Patterson, Aida M. Andrs, Evan E. Eichler, Montgomery Slatkin, David Reich, Janet Kelso, and Svante Pääbo. A high-coverage genome sequence from an archaic Denisovan individual. *Science*, 338 (6104):222–226, 2012.

[4] Nick Patterson, Swapan Mallick, and David Reich. *D*-statistics and interbreeding between archaic and modern humans. Supplementary Material 11 of Meyer et al. [3], 2012.

[5] Alan R. Rogers and Ryan J. Bohlender. Bias in estimators of archaic admixture. *Theoretical Population Biology*, 100:63–78, March 2015. ISSN 0040-5809.

[6] Paul R Staab, Sha Zhu, Dirk Metzler, and Gerton Lunter. Scrm: Efficiently simulating long sequences using the approximated coalescent with recombination. *Bioinformatics*, 31(10):1680–1682, 2015.
